# Can whole-lake algal biomass be captured by one-dimensional modeling approaches? An exploration using ‘Lake2D’

**DOI:** 10.1101/2025.05.02.651842

**Authors:** Hugo Harlin, Karl Larsson, Sebastian Diehl

## Abstract

Basin morphometry can strongly affect lake-internal processes relevant for productivity, such as turbulent mixing, photosynthetic energy acquisition, sedimentation, and nutrient recycling. Yet, in both empirical and theoretical studies of whole-lake primary production, lake morphometry is often simplified to a single 1-dimensional measure – lake mean depth. Using the conceptual, process-based model ‘Lake2D’, we addressed the question: To what extent can pelagic and benthic producer dynamics, integrated over a lake basin, be captured by approaches that use mean depth as the only morphometrical variable? We created two models of algal biomass dynamics in a radially symmetric, cone-shaped lake – one preserving the lake’s vertical and radial dimensions and one preserving only the lake’s mean depth – and compared model predictions of algal biomass dynamics across a wide range of lake sizes, mixing conditions, water transparency, and nutrient content. Our analyses reveal that model predictions differ substantially but predictably in much of the investigated parameter space, and identifies the light environment set by lake depth, water clarity and pelagic nutrients, but also lake area, as main drivers of the differences. Most commonly, the model based on mean depth underestimates benthic algal biomass and overestimates pelagic algal biomass, the net effect on total biomass being a 5-50% underestimate in shallow lakes and a 5-20% overestimate in many deeper lakes. Since gross primary production (GPP) in our model scales with algal biomass, we believe that global estimates of lake GPP should be corrected for the systematic errors inflicted by the prevailing 1-dimensional approaches.

## Introduction

Lake productivity is governed by three types of physical drivers: climate, which determines the input of energy from the sun and the atmosphere; watershed characteristics, which determine material inputs from the catchment; and lake morphometry, i.e. the size, depth and shape of the lake basin, which mediates the impacts of the other drivers on internal processes such as turbulent mixing, photosynthetic energy acquisition, sedimentation, and nutrient recycling. Historically, limnologists have emphasized watershed drivers, in particular the input of mineral nutrients and colored dissolved organic carbon, as major controls of lake productivity (Likens, 1972; Thienemann, 1925; Williamson et al., 1999). More recently, climate change has spurred a growing interest in climatic drivers of lake productivity and its phenology (Adrian et al., 2009; Michelutti et al., 2005; O’Reilly et al., 2003). In contrast, basin morphometry has received less attention as a driver of these processes, and in a majority of comparative studies information on lake morphometry is typically restricted to a single 1-dimensional length measure, i.e. lake mean depth (Jeppesen et al., 1997; Nõges, 2009; Taranu & Gregory-Eaves, 2008).

Lake mean depth has been found to be negatively related to a whole suite of productivity measures such as pelagic total nutrient concentrations (Chow-Fraser, 1991; Håkanson, 2004; Nõges, 2009; Taranu & Gregory-Eaves, 2008), pelagic chlorophyll a concentration (Jeppesen et al., 1997; Liu et al., 2011; Morana et al., 2023; Nõges, 2009), whole-lake gross primary production and respiration (Klaus et al., 2022; Staehr et al., 2012) and methane emission (Li et al., 2020; Wik et al., 2016). Yet, the predictive power of lake mean depth varies tremendously between studies and response variables, and it has long been recognized that additional information on basin shape and vertical water column structure is needed to capture the influence of lake morphometry on productivity (Carpenter, 1983; Fee, 1979). This is particularly true when benthic productivity is considered, which depends primarily on the amount of light that reaches the lake bottom (Godwin et al., 2014; Puts et al., 2022a; Vadeboncoeur et al., 2001). Typically, lake mean depth is a poor predictor of benthic productivity at all levels of the food web (Downing et al., 1990; Duarte & Kalff, 1990; Hanson & Peters, 1984; Puts et al., 2022a). Only recently, more informative morphometric descriptors have been used to predict the contribution of benthic habitats to whole-lake production, including the ratio of mean to maximum lake depth, which characterizes the depth distribution of the lake bottom, and the proportion of bottom area above the photosynthetic compensation depth (Norman et al., 2022; Puts et al., 2022a; Seekell et al., 2021; Vadeboncoeur et al., 2008).

Process-based modeling represents another approach to the study of primary production where information on lake morphometry has frequently been restricted to the vertical dimension. While production modules have been linked to 3-dimensional hydrodynamical models in the description of real lakes (Bruggeman & Bolding, 2014; Soares & do Carmo Calijuri, 2021), conceptual models with the primary purpose of gaining general process understanding have largely been limited to the description of pelagic producers in a 1-dimensional water column. These models predict that pelagic biomass should be maximized at moderate water column depths and turbulences (Diehl, 2002; Huisman & Sommeijer, 2002; Huisman & Weissing, 1995; Jäger et al., 2010), but should go extinct under three distinct circumstances: when high turbulence combines with a deep water column (causing excessive light limitation); at very low turbulence (causing excessive sinking losses and/or nutrient limitation); or in very shallow water columns (causing excessive sinking losses) (Diehl, 2002; Huisman et al., 1999; Jäger et al., 2010; Yoshiyama et al., 2009). It is by no means obvious how these interactive effects of depth and turbulence play out in the setting of a lake basin with spatially varying water column depth, and whether a 1-dimensional model could capture the average dynamics observed across the full 3-dimensional space.

A single 1-dimensional depth measure provides an even less suitable physical setting for process-based modeling of benthic primary producers. Yet, the few conceptual models that address benthic primary production are either zero-dimensional or assume a 1-dimensional water column of uniform depth (Jäger and Diehl, 2014; Klausmeier and Litchman, 2001; Scheffer, 1998; Vasconcelos et al., 2018; Yoshiyama et al., 2009; but see Genkai-Kato et al., 2012). An obvious weakness of these models is that they predict extinction of benthic primary producers when the amount of light that reaches the uniform bottom falls below the compensation depth (Vasconcelos et al., 2018). Yet, the same producers would always survive in shallower parts of a 3-dimensional lake basin even if the mean depth of the lake exceeds the compensation depth. Conversely, while a 1-dimensional model of a shallower water column predicts dominant benthic producers to block the transport of nutrients from the sediment to pelagic producers (Jäger & Diehl, 2014), it seems plausible that a corresponding 3-dimensional model with the same mean depth might predict a sustained nutrient flux to the pelagic from unvegetated sediments in deeper parts of the lake.

The above empirical and theoretical considerations beg the question: To what extent can pelagic and benthic producer dynamics, integrated over the bottom topography of a given lake basin, be captured by approaches that use lake mean depth as the only morphometrical input variable? And how does the answer to this question depend on other physical drivers such as lake size and depth, water clarity, nutrient content, and turbulent mixing? To our knowledge, this has never been investigated from a purely conceptual perspective, using a process-based modelling approach. To address these questions, we compared the dynamic behaviors of two models of benthic and pelagic primary producers that are identical in all respects but lake bathymetry. Specifically, we formulated two model approximations of algal biomass dynamics in a radially symmetric, cone-shaped lake (Fig. 1): a 1-dimensional (1D) model preserving only the lake’s mean depth, and a 2-dimensional (2D) model preserving the lake’s vertical and radial horizontal dimensions (and thus its maximum depth and surface area) which we name ‘Lake2D’. We subsequently compared 1D and 2D model predictions of benthic and pelagic algal biomass dynamics across a wide range of lake sizes, mixing conditions, water clarity, and nutrient content. Our analyses reveal that 1D and 2D model predictions differ substantially in much of the investigated parameter space, and identifies the light environment set by lake depth, water clarity and pelagic nutrients as the main driver of the observed differences.

**Figure 1:**
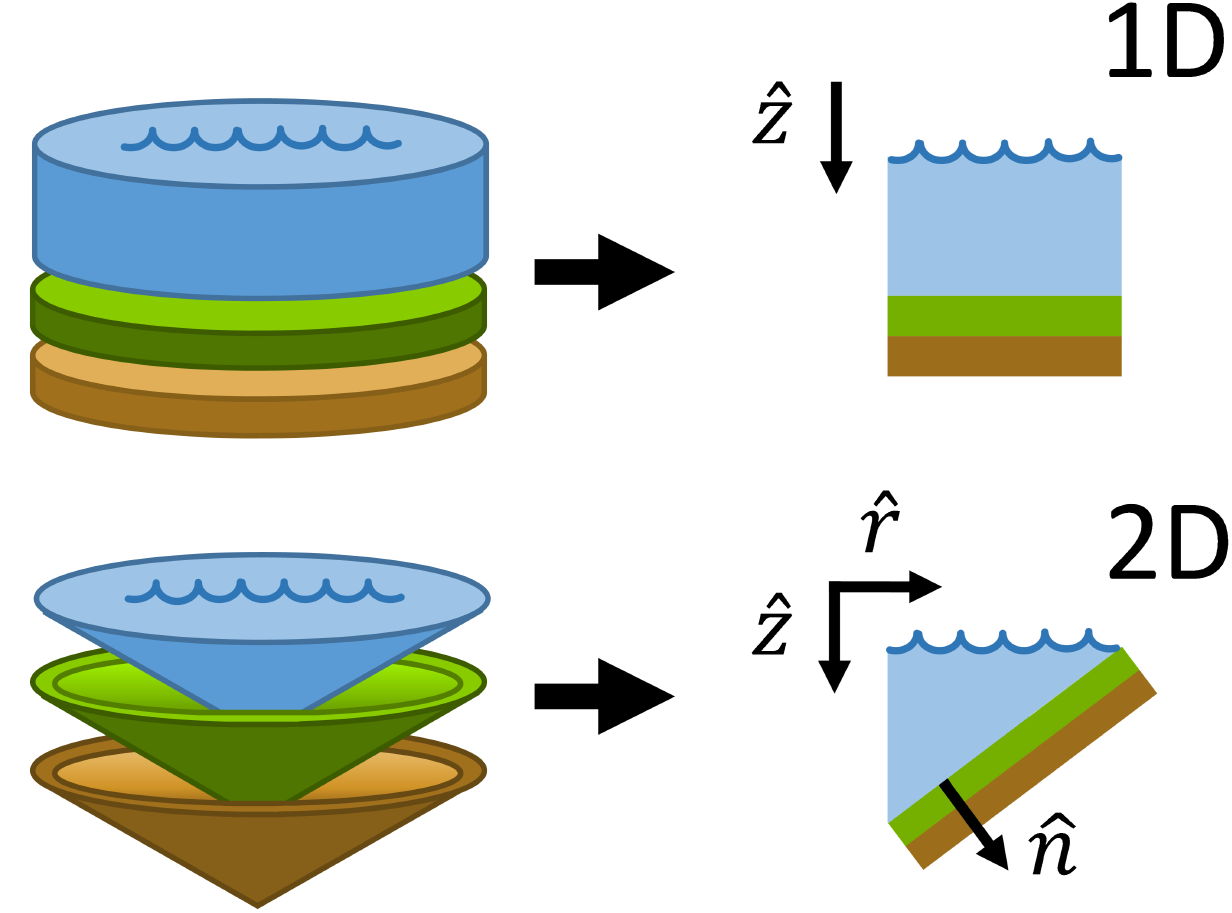
A hypothetical cylindrical and conical lake and their conceptual reduction to 1D and 2D lake models. The top row depicts the 1D model geometry, interpreted as a cross-section of a 3D cylinder-shaped lake, where the vertical dimension 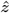 is the sole spatial variable. The bottom row depicts the 2D model geometry interpreted as a cross-section of a 3D cone-shaped lake that is radially symmetric, where the vertical and the radial 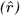 dimensions are the two spatial variables. The vector 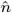 is the outward-facing normal vector to the lake bottom. Colors indicate the pelagic habitat (blue), a benthic algal layer (green), and a sediment layer (brown).

### Model description

Our 1D and 2D-models are derived from a full 3-dimensional model of algal biomass dynamics (Supplementary material A) using simplifying assumptions on lake geometry and symmetries on the problem. For the 2D-model Lake2D, we chose a radially symmetric cone-shaped lake because its simple topography facilitates visualization and interpretation of results, but also because its bottom profile is similar to the average hypsography of many real lakes (Håkanson, 1977; Klaus et al., 2022; Seekell et al., 2021; Vadeboncoeur et al., 2008). To isolate the impact of dimensionality in model comparisons, we used identical parameter values in 1D and 2D and furthermore assumed that benthic and pelagic algae have identical traits with respect to growth and metabolism. For the same reason, we assume that the intensities of horizontal and vertical mixing are independent of each other and do not vary in space. All variables and parameters are listed with their units in Table 1, using values from the literature (Jäger & Diehl, 2014; Jäger et al., 2010; Vasconcelos et al., 2018).

**Table 1:** State variables and parameters.

| Symbol | Unit | Value | Description |
| --- | --- | --- | --- |
| State variables |  |  |  |
| $A$ | $\text{mg C m}^{-3}$ | ... | Pelagic algal biomass concentration |
| $B$ | $\text{mg C m}^{-2}$ | ... | Benthic algal surface density |
| $I(z)$ | $\mu\text{mol photons m}^{-2}\text{s}^{-1}$ | ... | Light intensity in the lake water |
| $I(s)$ | $\mu\text{mol photons m}^{-2}\text{s}^{-1}$ | ... | Light intensity in the benthic algal layer |
| $N_B$ | $\text{mg P m}^{-3}$ | ... | Dissolved nutrient concentration in the benthic algal layer |
| $N_d$ | $\text{mg P m}^{-3}$ | ... | Dissolved nutrient concentration |
| $N_s$ | $\text{mg P m}^{-2}$ | ... | Benthic sedimented nutrient surface density |
| Variable environmental parameters |  |  |  |
| $d_r$ | $\text{m}^2 \text{ day}^{-1}$ | ... | Horizontal diffusion coefficient (2D-model) |
| $d_R$ | $\text{day}^{-1}$ | [0.001, 10] | Horizontal transport rate |
| $d_z$ | $\text{m}^2 \text{ day}^{-1}$ | ... | Vertical diffusion coefficient |
| $d_Z$ | $\text{day}^{-1}$ | [0.001, 100] | Vertical transport rate |
| $k_{bg}$ | $\text{m}^{-1}$ | 0.3, 1, 2 | Background light attenuation coefficient |
| $N_{\text{area}}$ | $\text{mg P m}^{-2}$ | 100, 300, 1000, 3000 | Total nutrient content per unit surface area |
| $r$ | m | ... | Distance from lake center in radial direction |
| $R$ | m | ... | Lake surface radius |
| $z$ | m | ... | Vertical distance from the water surface |
| $z_m$ | m | 1, 3, 10, 30 | Lake mean depth (2D-model), or max depth (1D-model) |
| Constant parameters |  |  |  |
| $d_b$ | $\text{m}^2 \text{ day}^{-1}$ | $10^{-6}$ | Benthic orthogonal diffusion coefficient (2D-model) |
| $d_{N_B}$ | $\text{m}^2 \text{ day}^{-1}$ | $1.7 \times 10^{-4}$ | Benthic perpendicular diffusion coefficient |
| $f$ | $\text{mg C mg}^{-1}$ | 0.1 | Benthic carbon mass to wet weight ratio |
| $G_{\text{max}}$ | $\text{day}^{-1}$ | 1.5 | Maximum specific growth rate |
| $I_0$ | $\mu\text{mol photons m}^{-2}\text{s}^{-1}$ | 300 | Light intensity at lake surface |
| $k$ | $\text{m}^2 \text{ mg C}^{-1}$ | $3 \times 10^{-4}$ | Light attenuation coefficient of algal biomass |
| $L$ | m | ... | Thickness of the benthic algal layer |
| $M$ | $\text{mg P m}^{-3}$ | 4 | Half saturation constant of nutrient-dependent algal production |
| $H$ | $\mu\text{mol photons m}^{-2}\text{s}^{-1}$ | 60 | Half-saturation constant of light-dependent algal production |
| $l_d$ | $\text{day}^{-1}$ | 0.02 | Mortality loss rate of algae |
| $l_r$ | $\text{day}^{-1}$ | 0.08 | Respiration loss rate of algae |
| $q$ | $\text{mg P mg C}^{-1}$ | 0.0015 | Algal stoichiometric P:C ratio |
| $\rho_w$ | $\text{mg m}^{-3}$ | $997 \times 10^6$ | Density of water |
| $r_s$ | $\text{day}^{-1}$ | 0.02 | Mineralization rate of sedimented nutrients |
| $v$ | $\text{m day}^{-1}$ | 0.1 | Algal sinking speed |

### One-dimensional model

The 1D-model assumes a horizontally well-mixed cylindrical water column with depth *z*_*m*_ and radius *R* corresponding to the mean depth and radius, respectively, of the cone-shaped 3D lake. It describes the dynamics of four state variables: the concentrations of pelagic algal biomass *A* and a potentially growth limiting inorganic nutrient *N*_d_ in the lake water, and the areal densities of sedimented particular nutrient *N*_s_ at the (flat) lake bottom and of benthic algal biomass *B* at the sediment-water interface. Benthic and pelagic algal biomass is expressed in units of carbon, and the dissolved and sedimented nutrient is assumed to be phosphorus. The rates of change of these state variables are governed by four differential equations

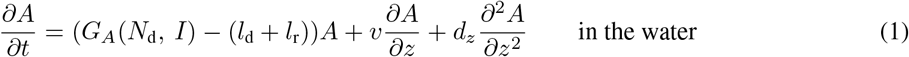

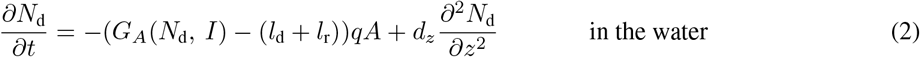

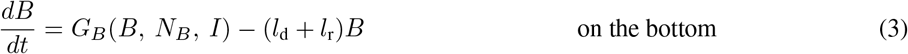

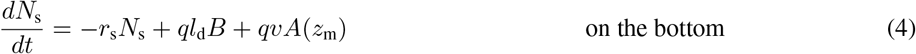

where the vertical gradient in the (potentially growth-limiting) intensity of light *I* in the water column follows Lambert-Beer’s law,

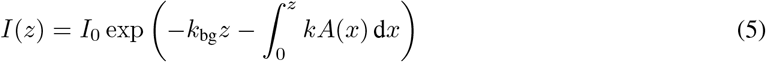

and the growth rates of pelagic and benthic algae, *G*_*A*_, and *G*_*B*_, are given by

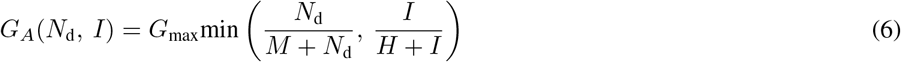

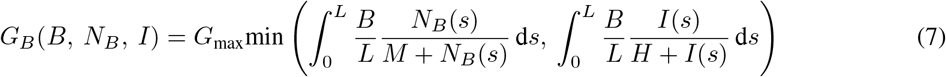

Equations 1–7 are explained in detail below. Parameter values and units are listed in Table 1.

### Transport mechanisms

The model considers two spatial transport processes, turbulent mixing and sinking. Turbulent mixing of pelagic algae and dissolved nutrients is described by the vertical diffusion coefficient *d*_*z*_, which is constant in space and time. Algae and nutrients cannot diffuse across the upper and lower boundaries of the water column. In addition, pelagic algae sink at a constant speed *v*. Pelagic algae that reach the bottom die and their nutrients turn into sedimented particular nutrients. Together, these assumptions lead to the following boundary conditions at the surface (water depth *z* = 0) and the bottom (*z* = *z*_m_) of the lake:

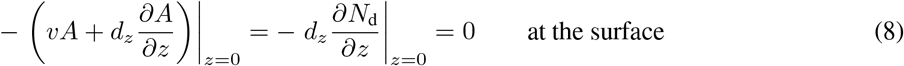

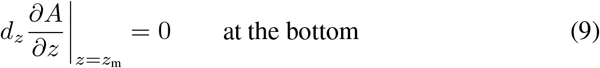

The bottom boundary condition for dissolved nutrients depends on several cross-boundary processes and is described further down.

### Pelagic algal dynamics

The growth of pelagic algae at a given depth *z* is either limited by the local light intensity, or the local dissolved nutrient concentration, as described by the minimum function of two Monod terms in (6) where *G*_max_ is the maximum specific growth rate and *H* and *M* are the half-saturation constants of light and nutrient limited growth, respectively. Light intensity *I* attenuates vertically in the water column according to (5), where pelagic algae contribute with specific attenuation coefficient *k* and background attenuation with coefficient *k*_bg_. The latter represents light attenuation by non-algal materials in the water and by water itself. In addition to sedimentation losses, pelagic biomass is lost through background respiration and mortality with fixed rates *l*_r_ and *l*_d_, respectively.

### Pelagic nutrient dynamics

The uptake and recycling of dissolved nutrients at a given depth are driven by local algal growth and losses. We assume that the stoichiometric phosphorus-to-carbon (P:C) ratio *q* of algal biomass is fixed. Consequently, the local uptake of dissolved phosphorus equals a fraction *q* of algal carbon production. Similarly, algal (carbon) biomass losses through mortality and respiration are accompanied by proportional phosphorus losses, which we assume to be instantly recycled in dissolved form.

### Sediment nutrient dynamics

Particular nutrients enter the sediment through two processes – sinking of pelagic algae and background mortality of benthic algae at the rate *l*_d_. We assume that remineralization of sedimented nutrients is a first order process at the rate *r*_s_ and that remineralized nutrients are released into the water directly above the bottom (see (14)).

### Benthic algal dynamics

Our assumption that benthic and pelagic algae have identical traits implies that benthic and pelagic algae have identical maximum specific growth rates, identical half saturation constants for light and nutrient dependent growth, identical light attenuation coefficients and elemental P:C ratios, as well as identical background respiration and mortality rates.

To calculate their growth rate, we assume that benthic algae form a homogeneous layer of thickness *L*. The value of *L* depends on benthic algal biomass according to the equation

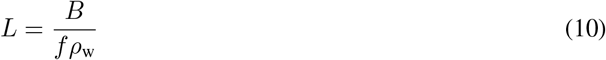

where *f* is the ratio of benthic algal carbon to wet mass and *ρ*_w_ is the specific density of algal wet mass assumed to be identical to the density of water. Light in the benthic algal layer is attenuated according to Lambert-Beer’s law. Assuming that non-algal light attenuation is negligible, light-limited benthic algal growth can then be integrated over the benthic algal layer, which has the explicit solution (Huisman & Weissing, 1994)

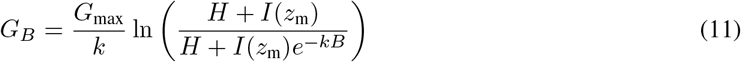

where *I*(*z*_m_) is the local light intensity at the bottom of the water column.

Nutrient-limited benthic algal growth is modeled analogously by integrating over the vertical profile of dissolved nutrients in the benthic algal layer. We assume that dissolved nutrients from immediately above the lake bottom diffuse into the benthic algal layer while simultaneously being consumed by benthic algae. These two processes are modeled with the differential equation

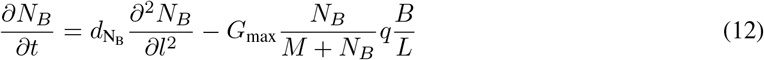

where *N*_*B*_ is the concentration of dissolved nutrient in the benthic algal layer, 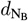 is the diffusion coefficient in the benthic algal layer, *l* ∈ [0, *L*] denotes the vertical position inside the benthic algal layer, and *q* is the P:C ratio of benthic biomass. We assume that these processes are fast compared to other processes, and therefore treat (12) on a separate timescale. For each time *t*, we approximate the steady state solution of (12) by linearizing the benthic algal consumption term and solving the resulting equation analytically. We further assume that the nutrient concentration at the surface of the benthic algal layer equals the concentration at the bottom of the water column and that nutrients do not diffuse out of the bottom of the benthic algal layer, yielding the boundary conditions

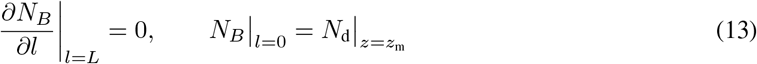

We then integrate nutrient limited growth over the benthic algal layer (first integral in (7)), using the approximated solution of (12) for each time step. In a final step, benthic algal growth per unit area is calculated as the minimum of nutrient and light-dependent growth (7). This approach can sometimes overestimate benthic algal growth, since it is possible that growth switches from nutrient to light limitation somewhere inside the benthic algal layer. Note that *N*_*B*_ is a dummy variable that is only used in the calculation of nutrient-limited benthic algal growth and therefore does not contribute to the nutrient mass balance of the system.

### Nutrient fluxes at the sediment-water interface

We assume that nutrients lost through benthic algal respiration are released in dissolved form into the water directly above, which is the same water from which the nutrients are drawn that sustain benthic algal growth. Together with the release of remineralized particular nutrients from the sediment, this yields the following boundary condition for dissolved nutrients at the bottom of the water column

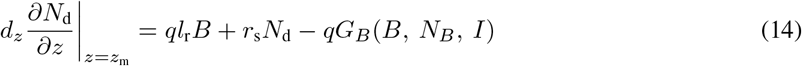

### Two-dimensional model

The formulation of the model Lake2D is identical to the 1D-model with two distinctions: it incorporates a cone-shaped lake morphometry with a linearly sloping bottom (with the same mean depth *z*_m_ and radius *R* as the 1D-model) and it includes horizontal transport processes. We also assume radial symmetry such that, for any distance from the center vertical axis of the lake, all state variables are well mixed in the tangential direction. Two spatial variables are then sufficient to describe spatial dynamics: a radial direction 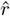 and a vertical direction 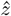 (Fig. 1). We made the simplifying assumption of radial symmetry because we wanted to investigate the effects of a sloping lake bottom topography and the possibility of horizontal gradients in pelagic algae and nutrients on system dynamics, while keeping model complexity to a minimum and enabling the comprehensive illustration of results in 2D (cf. Fig. 4-6). The computationally expensive calculation of the full 3D dynamics of the system is not necessary for addressing the research questions posed in this paper.

### Horizontal transport processes

In the pelagic habitat, horizontal turbulent mixing in the radial direction is modelled (in cylindrical coordinates) by adding the terms 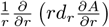 to (1) and 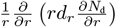 to (2), where *r* is the distance from the lake center and *d*_*r*_ is the horizontal diffusion coefficient in radial direction. We also enabled horizontal spread of benthic algae and sedimented nutrients by adding a very small radial diffusion term to (3) and (4). See Supplementary material B for details.

### Model implementation

To save computation time, we implemented Lake2D on a half cross-section of the lake (Fig. 1). The full 3D cone-shaped geometry can be retrieved by rotating this triangular cross-section around the center axis of the lake. This approach necessitates a reflective boundary condition for pelagic algae and dissolved nutrients along the left boundary of the triangular cross-section, corresponding to the center of the lake:

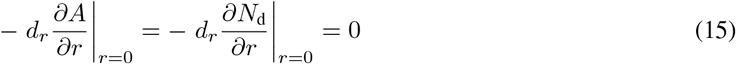

### Equivalence of models

The 1D and 2D-models should be regarded as three-dimensional systems that have been simplified to fewer dimensions for calculation purposes. They are equivalent in the following ways:

- The models are derived from the same physical three-dimensional system (Supplementary material A) with the same parameters with identical values (*l*_r_, *l*_d_, *H, M*, *G*_*A*_, *G*_*B*_, *k*_bg_, etc.).
- The models have the same mean depth (*z*_m_), surface area (*πR*^2^), and volume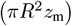.

### Nutrient conservation

The absence of internal nutrient sources and the closed boundary conditions imply that a modeled lake’s total nutrient content *N*_area_, defined as total phosphorus content per unit lake surface area, is determined by the initial conditions and remains constant over time. *N*_area_ is defined for the 1D-model as

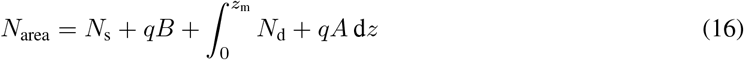

and for the 2D-model as

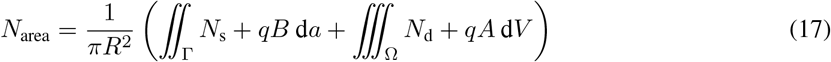

where Γ is the surface area, and Ω is the lake volume.

## Methods

### Design of numerical experiments

To assess how 1D and 2D-model predictions compare with each other, we performed extensive numerical simulations focused on six environmental parameters that govern the underwater light conditions and/or influence the formation of spatial gradients. Specifically, we varied background turbidity (*k*_bg_) and lake mean depth (*z*_m_), which directly affect light conditions, as well as the total nutrient content per surface area (*N*_area_), which indirectly affects light conditions by establishing an upper limit on potential shading by pelagic algal biomass. We furthermore varied lake area (by varying the radius *R*) and the horizontal and vertical diffusion coefficients (*d*_*r*_, *d*_*z*_) which, together with lake mean depth, influence the formation of horizontal and vertical gradients. To reduce the dimensionality of the problem, we combined the horizontal diffusion coefficient with the lake radius into a single, aggregate parameter

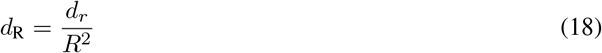

called the horizontal mixing rate. We similarly combined the vertical diffusion coefficient with the lake mean depth into a single parameter

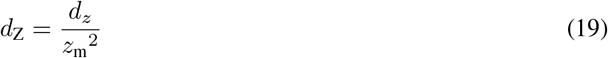

called the vertical mixing rate. The effects of varying *R* and *d*_*r*_ are fully captured by the horizontal mixing rate *d*_R_, such that a given change in the value of *d*_R_ can be interpreted as a change in either lake surface area or the intensity of horizontal mixing. Since the horizontal dimension of most real lakes exceeds their vertical dimension by several orders of magnitude, and the intensity of turbulent mixing (as captured by the turbulent diffusion coefficients *d*_*r*_ and *d*_*z*_) is typically several orders of magnitudes higher in horizontal than in vertical direction (Peeters et al., 1996), we prefer the former interpretation of the horizontal mixing rate *d*_R_ as being synonymous to an inverse measure of lake area. Similarly, a given change in the value of *d*_Z_ can be interpreted as a change in either lake mean depth or the intensity of vertical mixing. Yet, since pelagic algae sink at a constant speed, *d*_Z_ does not capture all effects of lake depth, and *z*_m_ needs to be treated as an independent parameter. Together, *d*_R_ and *d*_Z_ constitute a two-dimensional slice of parameter space that we hereafter refer to as the ‘mixing space’.

Collectively, lake mean depth *z*_m_, background attenuation coefficient *k*_bg_, areal nutrient content *N*_area_, and the mixing space define a 5-dimensional parameter space, which we numerically explored with both the 1D and 2D-model in a fully factorial design, where the parameters *k*_bg_, *z*_m_, and *N*_area_ were varied over ranges that are representative of the vast majority of the world’s lakes (Cael & Seekell, 2022; Cael et al., 2017; Chen et al., 2015; Sobek et al., 2007). Specifically, we performed 19200 numerical simulations for the Lake2D model resulting from the combination of four values of *N*_area_ (100, 300, 1000, and 3000 mgP m^*−*2^), four values of *z*_m_ (1, 3, 10, and 30 m), three values of *k*_bg_ (0.3, 1, and 2 m^*−*1^), and a 20 *×* 20 factorial mixing space, where the horizontal and vertical mixing rates *d*_R_ and *d*_Z_ were varied on a logarithmic scale over the intervals [0.001, 10] day^*−*1^ and [0.001, 100] day^*−*1^, respectively. The corresponding number of 1D-model simulations was 960 (because *d*_R_ is not part of that model).

### Execution of numerical experiments

To increase computational efficiency, the models were first non-dimensionalized in space and time (see Supplementary material B). The models were implemented in Matlab using a finite volume method and the built-in function ODE15s was used to integrate in time. A discussion of the finite volume method’s strengths is provided in Supplementary material B. A diagonally cut grid of 20×20 equally spaced grid cells was frequently sufficient to achieve accurate results. Under certain environmental conditions, steep spatial gradients could arise at specific depths or distances from the lake center. In such cases, the local grid resolution near the steep gradients was increased as necessary to maintain accuracy. All simulations were run to steady state, and the same initial condition was used for all simulations; 98% of total nutrients in dissolved form, 1% in pelagic algae, and the remaining 1% in benthic algae, all evenly distributed across the lake’s volume and bottom, respectively.

### Model output and visualization

We focus our analyses on a comparison of 1D and 2D-model output at the whole-lake scale. For each individual model run, we therefore integrated steady-state pelagic and benthic algal biomass over the entire lake and expressed it as average biomass per unit of lake volume and surface area, respectively. To visualize the results in the complex 5-dimensional parameter space, we proceeded as follows. For each combination of *z*_m_, *k*_bg_, and *N*_area_ values, we first produced heat maps of the steady-state pelagic and benthic algal biomasses in *d*_R_ x *d*_Z_ mixing space for model Lake2D and in *d*_Z_ mixing space for the corresponding 1D-model (see Fig. 2 a-f, j-o). We then calculated 1D:2D whole lake biomass quotas for each mixing space pair by dividing the biomass obtained in 1D with the corresponding biomass obtained in 2D for each value of the horizontal mixing rate *d*_R_, yielding two-dimensional heat maps of 1D:2D biomass quotas in *d*_R_x*d*_Z_ mixing space (see Fig. 2 g-i, p-r). In a final step, we organized the resulting 1D:2D biomass quota heat maps in a multi-panel figure in which heat maps with different background turbidities *k*_bg_ are nested within columns of total nutrient content *N*_area_ and rows of lake mean depth *z*_m_ (Fig. 3).

**Figure 2:**
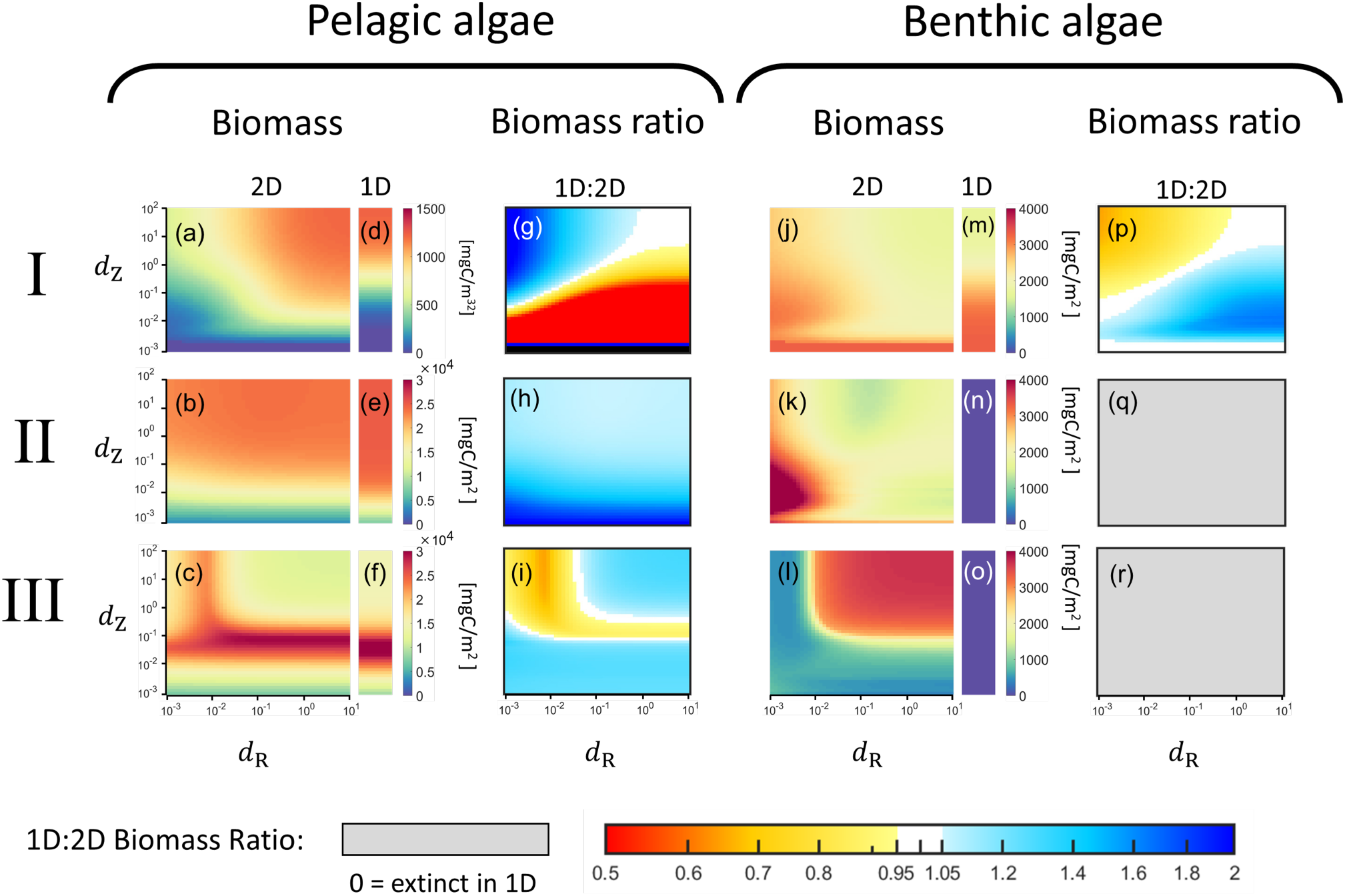
Heat map plots of pelagic and benthic algae in vertical (d_Z_) and horizontal (d_R_) mixing space, where each row corresponds to one of the three highlighted patterns I, II, and III in Fig. 3. Panels (a)-(f) display the lake-wide average surface densities of pelagic algae in mg C m^−2^ for the 2D-model and the 1D-model, respectively. Panels (g)-(i) display the corresponding 1D:2D pelagic algal biomass quotas. Panels (j)-(o) display the lake-wide average surface densities of benthic algae in mg C m^−2^ for the 2D model and 1D model, respectively. Panels (p)-(r) display the 1D:2D biomass quotas of benthic algae. Heat maps of biomass use the color scales displayed in vertical bars. Heat maps of 1D:2D quotas use the color scale displayed at the bottom.

**Figure 3:**
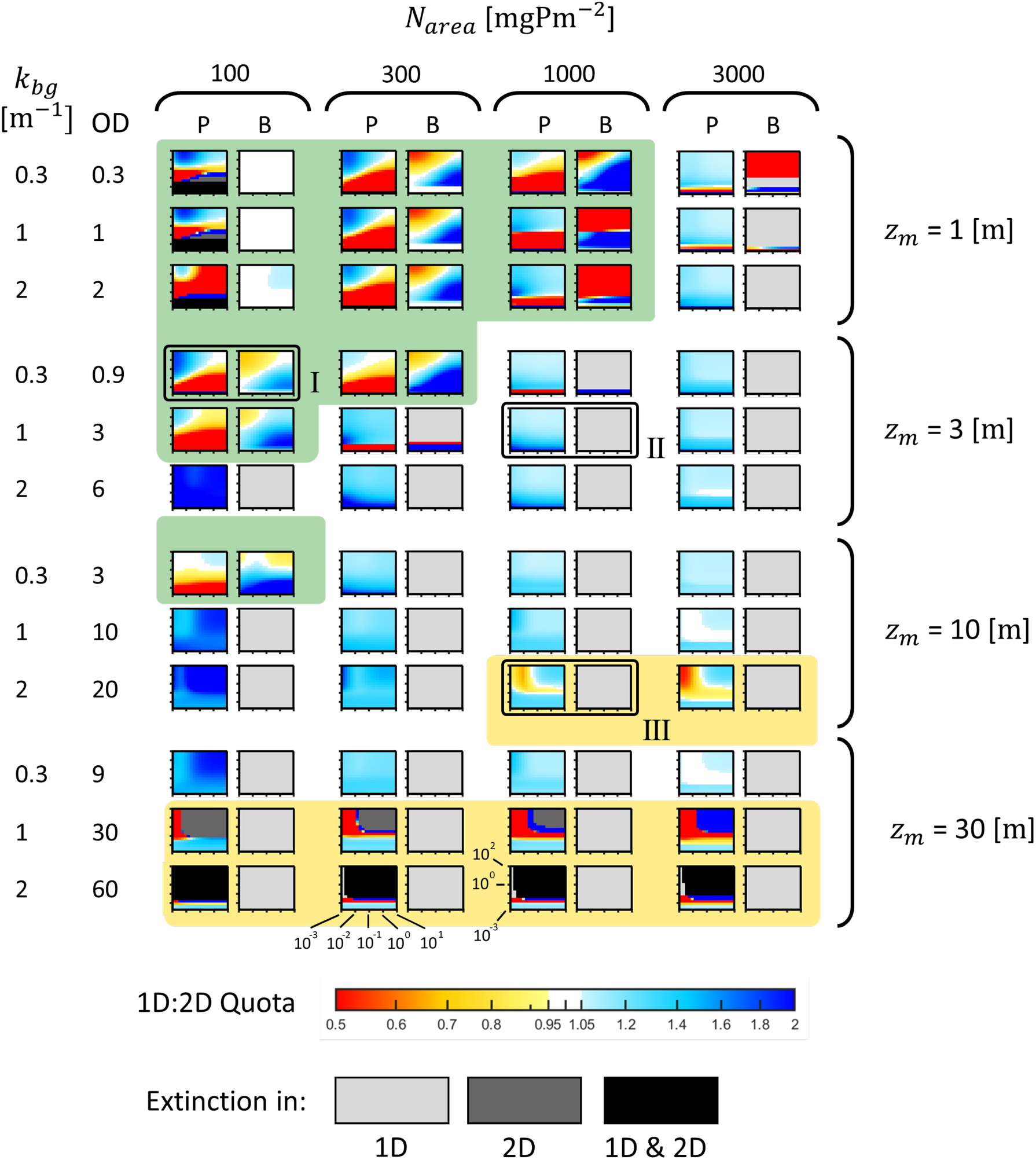
Heat map pairs of 1D:2D quotas of lake-wide total pelagic (P) and benthic (B) algal biomass for varying background turbidity k_bg_, total areal nutrient content N_area_, mean depth z_m_, and diffusive mixing rates d_R_ and d_Z_. The horizontal diffusive mixing rate d_R_ is varied along the horizontal axis of each panel in the range [0.001,10] day^−1^, and the vertical diffusive mixing rate d_Z_ is varied along the vertical axis of each panel in the range [0.001,100] day^−1^. Both axes are plotted on a log10 scale. Optical depth (OD = k_bg_z_m_) is shown next to each lake type’s background turbidity Each pixel in a panel is the quota of a 1D and 2D simulation pair, where the color represents the value of total pelagic or benthic biomass at steady state in 1D, divided by the same quantity in a corresponding 2D simulation with identical vertical mixing rate d_Z_. A reddish color corresponds to more algae in the 2D simulation, and a blue color corresponds to more algae in the 1D simulation. Quotas that deviate less than 0.05 from 1 are colored white, see the colorbar at the bottom of the figure. Grayscale is used to indicate extinction: Light gray indicates that algae have gone extinct in the 1D simulation but not in the 2D simulation, dark gray indicates that algae have gone extinct in 2D but not in 1D, and black indicates that algae have gone extinct in both 2D and in 1D. The three framed heat map pairs have been selected to highlight the three major patterns I, II, and III in mixing space. Patterns I and III are highlighted with a green and yellow background, respectively, to visually distinguish them from pattern II.

Inspection of the 1D:2D biomass quota heat maps in Fig. 3 revealed the existence of three qualitatively distinct patterns by which the five investigated environmental parameters affect the accuracy with which the 1D-model captures the spatially integrated algal biomass dynamics in our idealized cone-shaped lake. A deeper understanding of these patterns requires a closer examination of the within-lake spatial distributions of nutrients and algae under different mixing scenarios. To facilitate this, we selected one representative example of each of the three patterns (patterns I-III in Fig. 2) for which we show individual 1D and 2D simulation output from the four corners in mixing space, representing all four possible combinations of high and low vertical and horizontal mixing rates. The shown output includes the steady-state spatial distributions of pelagic and benthic algae, pelagic and sedimented nutrients, the relative partitioning of total nutrients among these four state variables, and the spatial distribution of the specific growth rate of pelagic algae (Fig. 4-6).

**Figure 4:**
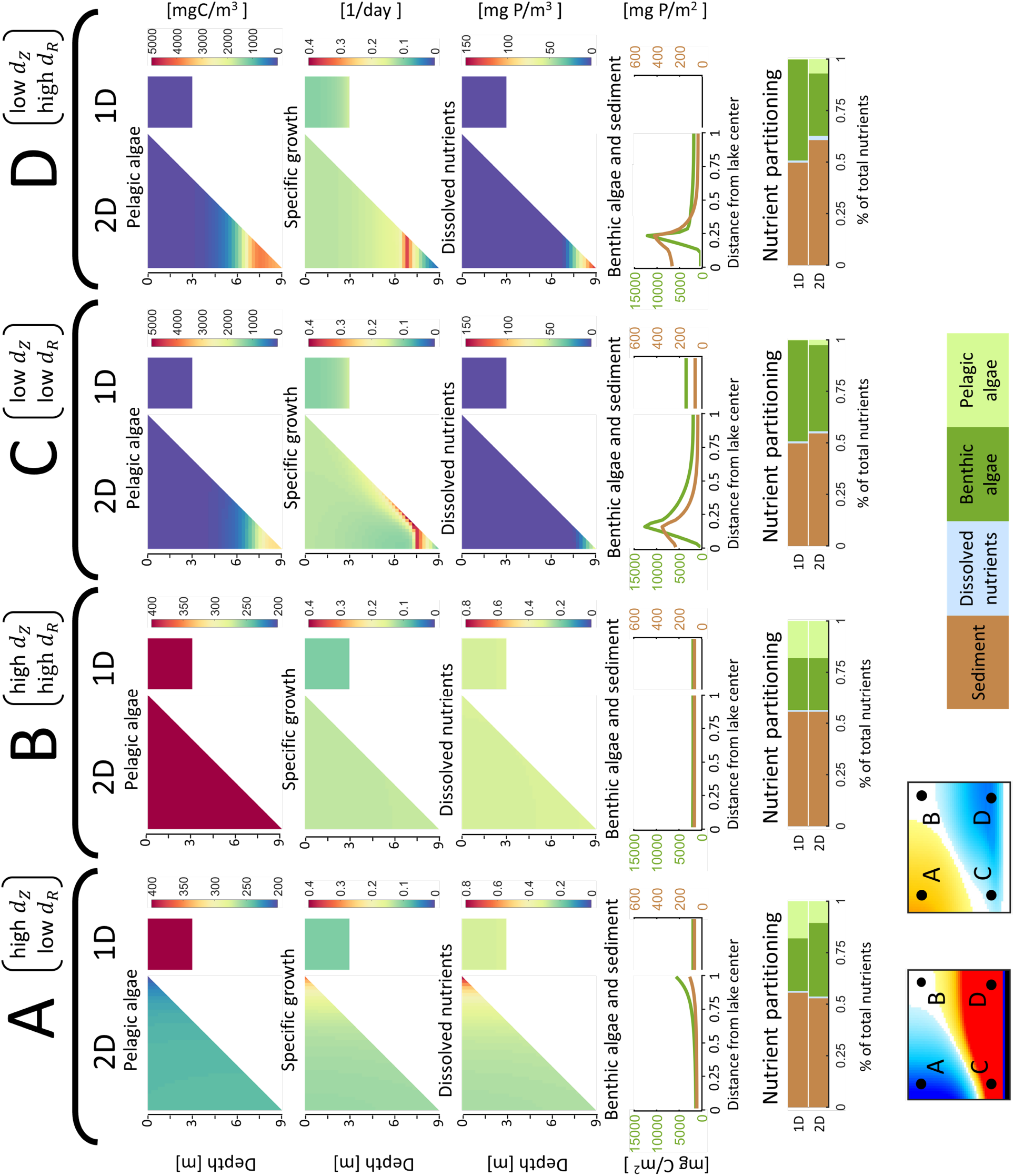
Four example simulations visualizing 2D- and 1D-model output that yields pattern I in Fig. 3. Each of columns A-D displays one pair of 2D- and 1D-simulations selected from the four corners of the vertical and horizontal mixing space (insets at bottom left), with the 2D-model output to the left and the 1D-model output to the right in each column. In panel rows 1-4, the vertical left edge of the 2D-panels corresponds to the center of the lake, and the x-axis is the scaled distance from the lake center. Since the horizontal axis contains no information in the 1D-model, the width of 1D-panels has been compressed. The **first row** of panels shows the pelagic algal biomass concentration (mg C m^−3^), the **second row** the specific growth rate of pelagic algae (day^−1^), and the **third row** the dissolved pelagic nutrient concentration (mg P m^−3^). The **fourth row** shows the standing stocks of benthic algal biomass (green lines, mg C m^−2^) and sedimented nutrients (brown lines, mg P m^−2^) per lake surface area. The bottom row of horizontal bar plots shows the partitioning of the system’s total nutrient content among the four state variables, with sediment in brown, dissolved nutrients in blue, and benthic and pelagic algae in dark and light green, respectively. Each pair of bars shows 1D-output on top and 2D-output below. Since the system’s total nutrient content is fixed, bars can be directly compared across all panels.

### Model runs including a mixed surface layer

Our standard model makes the simplifying assumption that vertical mixing intensity is homogeneous over lake depth. A large fraction of the world’s lakes is, however, seasonally or permanently stratified such that vertical mixing is high in a well-mixed surface layer, which is separated from less intensely mixed waters below. To explore the robustness of the results from the standard model against relaxation of the assumption of homogeneous vertical mixing, we re-ran all 1D- and 2D-simulations with two additional model variants for which we assumed a well-mixed surface layer (*d*_Z_ = 280) of 1.5 m and 4.5 m depth, respectively, while keeping horizontal mixing and mixing below the mixed surface layer identical to the standard model. We evaluated the robustness of the results from the standard model by comparing them to the output from corresponding stratified models across the entire 5-dimensional environmental parameter space.

## Results

### 1D:2D-model comparisons - overarching observations

Fig. 3 synthesizes the results of our comparison of the 1D- and 2D-models in the investigated 5-dimensional environmental parameter space. Inspection of Fig. 3 yields several overarching observations and general insights:

- The 1D-model very rarely gives similar results as the 2D-model (indicated by the rarity of white regions in the panels of Fig. 3). Thus, 1D and 2D simulation outcomes at the whole lake scale differ as a rule, the most common deviations being that the 1D-model underestimates the biomass of benthic algae (indicated by a dominance of light gray and reddish regions in the benthic panels of Fig. 3) and overestimates the biomass of pelagic algae (indicated by a dominance of blueish regions in the pelagic panels of Fig. 3). Yet, the reverse inaccuracies do also occur under some circumstances (indicated by blueish regions in the benthic panels and reddish regions in the pelagic panels of Fig. 3, respectively).
- Benthic algae always persist where a lake bottom is not aphotic. Consequently, benthic algae cannot go extinct in sufficiently shallow parts of a 2D-model system, but will go extinct in the 1D-model when the uniform lake depth exceeds the compensation depth (Fig. 2n, and Fig. 3, light gray panels).
- At very low vertical mixing rate *d*_*z*_, pelagic algae always decline with a further decrease in mixing (Fig. 2 a-f) and eventually go extinct in both 1D- and 2D-models, because turbulent mixing and local production cannot counter sinking losses. In most panels of Fig. 2 and 3, the extinction regions are outside of the depicted mixing space, except for the shallowest and most nutrient-poor systems where sinking losses are highest and production is lowest (black areas in upper left panels, Fig. 3).
- In very dark and/or deep lakes, the majority of the lake volume is aphotic. High vertical mixing therefore causes severe light limitation of pelagic algae and can drive them to extinction, unless horizontal mixing is so low that shallow regions can serve as a refuge in the 2D-model (e.g. dark gray vs. red areas in second-last row of panels from the bottom, Fig. 3)
- The deviations of the 1D-from the 2D-model output can be categorized into three qualitatively different patterns in horizontal and vertical mixing space (subsequently called patterns I-III), which depend on how the remaining three parameters lake mean depth, background attenuation, and nutrient supply together shape the light and nutrient environment. Below we describe each of patterns I-III and explore the underlying mechanisms by comparing within-lake spatial distributions of processes and state variables in selected mixing scenarios. In these descriptions, because light attenuates with depth in proportion to the product of depth times the light attenuation coefficient, we will frequently use optical depth as a dimensionless measure of the abiotic light conditions in the lake water, defined

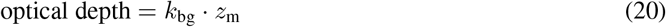

### Major patterns in mixing space

#### Pattern I: Diagonal gradients in biomass quotas in mixing space

This pattern occurs in shallow, high light (optical depth *≤* 3), low nutrient environments (Fig. 2, top row of panels, and Fig. 3, panels on green background), where a large fraction of the nutrient pool is stored in the sediment, benthic biomass exceeds pelagic biomass, and algal growth tends to be nutrient limited everywhere under most mixing scenarios. Under such conditions, pelagic algae benefit from both vertical and horizontal mixing (Fig. 2a) because mixing counteracts sinking and redistributes remineralized sediment nutrients both up in the water column and away from near-bottom areas where sinking losses are highest, reducing the dissolved nutrients available to the benthic algae. Benthic algae show the opposite diagonal pattern (Fig. 2j), i.e. benefit from low vertical and horizontal mixing and the resulting high pelagic sinking losses, which transport limiting nutrients to the bottom layer.

Because the 1D-model only allows for gradients in the vertical mixing direction, the 1D:2D comparisons generate diagonal patterns in mixing space, where the 1D-model typically either over- or underestimates benthic and pelagic biomass in a complementary way (Fig. 2g, p). These inaccuracies stem from the 1D-model’s inability to capture the 2D-model’s uneven spatial distribution of biomass and nutrients, as illustrated by the following select mixing scenarios (Fig. 4):

At high vertical but low horizontal mixing (Fig. 4, Scenario A), slow but steady horizontal transport of both dissolved nutrients and pelagic algae to the shallowest areas (where nutrient uptake by benthic algae and pelagic algal sinking losses are highest) causes a build-up of benthic biomass and sediment in nearshore areas, which is subsequently maintained by local nutrient recycling (note the higher nearshore dissolved nutrient concentrations). Because these shallow areas take on a disproportionately large fraction of the 2D lake bottom (56% of the bottom of a cone-shaped lake is shallower than the mean lake depth), the 1D-model overestimates pelagic biomass and underestimates benthic biomass at the whole-lake scale (see ‘Nutrient partitioning’ in Fig. 4A).

At low vertical mixing (Fig. 4, Scenario C and D), slow upward transport promotes the concentration of pelagic algae and nutrients near the lake bottom. In the 1D-model, the relatively shallow lake depth and high concomitant sinking losses strongly limit pelagic biomass to the benefit of benthic algae. In contrast, in the 2D system, nutrients bound in sinking pelagic algae are funneled away from shallower lake bottoms (increasing benthic algal nutrient limitation) towards the deepest parts of the lake, where high dissolved nutrient concentrations fuel growth rates that support a thin but dense pelagic biomass maximum, shading out benthic algae below. This enables pelagic algae to reach significantly higher local – and lake-averaged – concentrations in 2D compared to 1D for the same areal nutrient content in the water. The 1D-model therefore underestimates pelagic biomass and overestimates benthic biomass at the whole-lake scale (see ‘Nutrient partitioning’ in Fig. 4A). Finally, at high vertical and horizontal mixing (Fig. 4, Scenario B), no spatial gradients in specific algal production can arise, because light is abundant everywhere and the strictly growth-limiting nutrient is homogeneously distributed. Moreover, average sinking losses in a cone-shaped and a cylinder-shaped lake of equal mean depth are identical when biomass is homogeneously distributed. Consequently, both models produce near-identical outcomes at the whole-lake scale in this scenario.

#### Pattern II: Vertical pelagic gradients in biomass quotas in mixing space

This pattern occurs in medium light environments, characterized either by an optical depth *≥* 3 and *<* 20 or by an optical depth *≤* 3 and relatively high nutrient supply that causes shading from pelagic algae (Fig. 3, panels on white background). In such environments, pelagic biomass typically greatly exceeds benthic biomass (Fig. 2, middle row of panels) and algal growth is nutrient limited near the surface, but light limited immediately below, in both the 2D- and 1D-models (indicated by specific growth rates 0.1/day below a narrow surface layer, 2nd panel row in Fig. 5). Sufficient upward mixing of recycled nutrients from deep waters and sediments is then required to produce surface growth rates that compensate for negative net growth at aphotic depths. Consequently, pelagic algae benefit from increased vertical mixing in both 2D and 1D (Fig. 2b, e). In contrast, horizontal mixing has only small effects on pelagic algal biomass in 2D (Fig. 2b), because weak horizontal mixing does not result in strong horizontal gradients in depth-integrated specific production.

**Figure 5:**
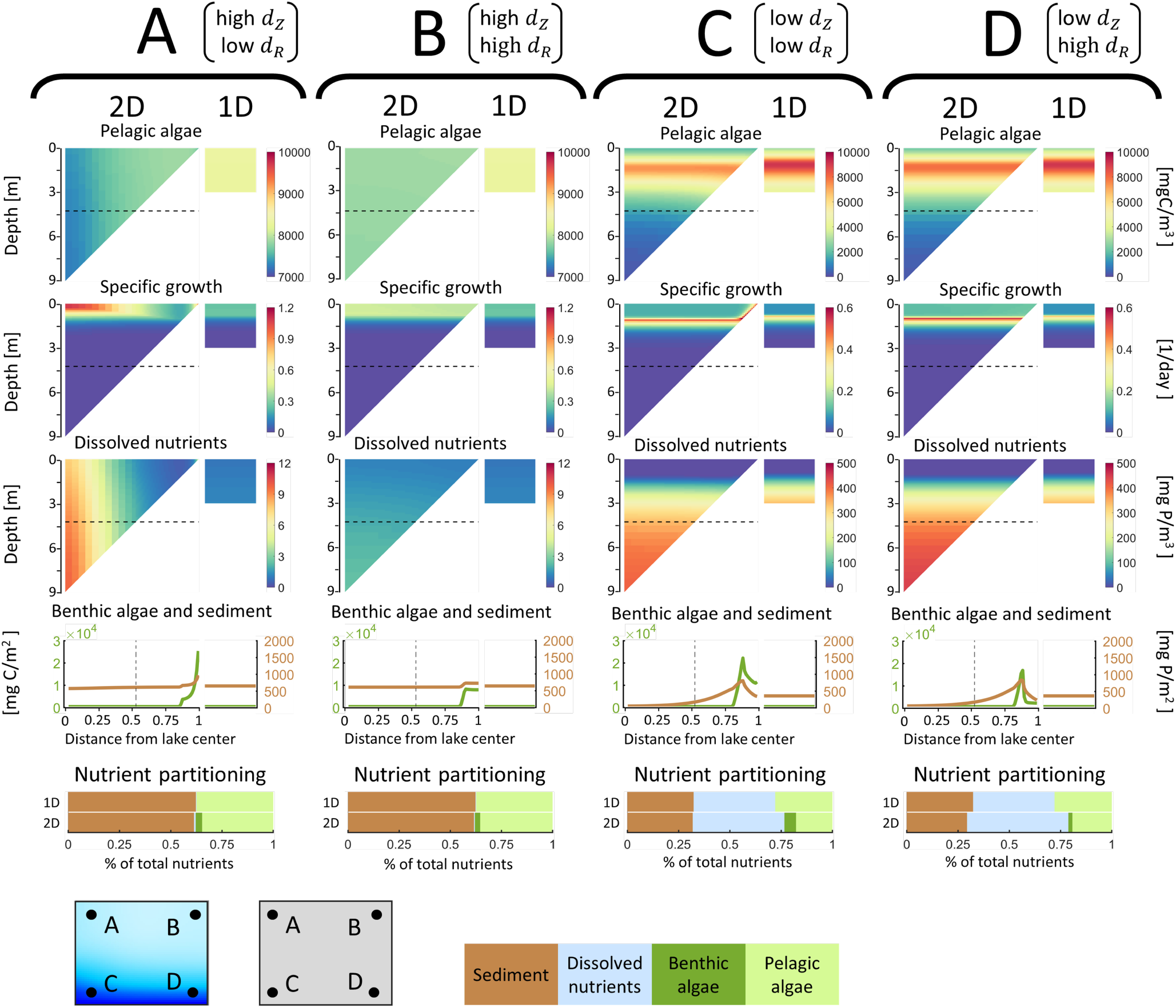
Four example simulations visualizing 1D- and 2D-model output that yields pattern II in Fig. 3. Horizontal broken lines in the first three panel rows mark the boundary of the euphotic depth, below which specific background losses exceed light-dependent production in a hypothetical system lacking pelagic algae. See caption of Fig. 4 for remaining details.

When horizontal mixing is low but vertical mixing is high (Fig. 5, Scenario A), surface growth rates are disproportionally high above the deepest parts of the lake, where the aphotic zone is the largest and thus generates both the highest population losses but also the highest recycling and upward transport of nutrients. Yet, depth-integrated specific net growth exhibits only weak horizontal gradients in spite of strong gradients in dissolved nutrients and depth-specific growth rate (Fig. 5, Scenario A). Pelagic biomass therefore reaches similar concentrations and shows an almost equally homogeneous spatial distribution as at high horizontal mixing (Fig. 5, Scenario A and B). Horizontal mixing has even less of an effect on pelagic biomass at low vertical mixing.

Under such conditions, pelagic algal sinking causes the majority of the nutrient pool to accumulate in dissolved form at strictly aphotic depths, where horizontal mixing is irrelevant to pelagic production because growth is light-limited (Fig. 5, Scenario C and D). This fraction of unused, dissolved nutrients is always lower in the 1D-model because a smaller fraction of its lake volume is aphotic (see ‘Nutrient partitioning’ in Fig. 5, Scenario C and D). The 1D-model therefore increasingly overestimates pelagic biomass with decreasing vertical mixing (Fig. 2h).

Finally, as long as vertical mixing is sufficient for pelagic algae to persist, the 1D-model always incorrectly predicts the extinction of benthic algae (Fig. 2n), because shading from pelagic algae creates an aphotic environment at the bottom of the 1D-lake, while benthic algae can persist in the shallow margins of the 2D-lake at biomass levels that are only weakly influenced by vertical and horizontal mixing (Fig. 2k, Fig. 5). In summary, in pattern II the 1D-model overestimates pelagic algal biomass – the more so, the lower the vertical mixing rate – and incorrectly predicts the extinction of benthic algae (see ‘Nutrient partitioning’ in Fig. 5).

#### Pattern III: L-shaped pelagic gradients in biomass quotas in mixing space

This pattern occurs in deep, low light environments characterized by an optical depth *≥* 20 (Fig. 2, bottom row of panels, and Fig. 3, panels on yellow background), where *>* 80 % of the lake volume is aphotic, benthic and pelagic production are light limited almost everywhere in the lake, and a large fraction of the nutrient pool is found in dissolved form at aphotic depths. Under such conditions, pelagic algae suffer from the combination of high horizontal and vertical mixing (Fig. 2c, Fig. 6 scenario B) because this mixing regime transports algae away from well-lit nearshore and surface areas to aphotic depths, and the upward transport of dissolved nutrients does not benefit pelagic algae much since they are predominantly light limited. In contrast, reduced mixing promotes the build-up and maintenance of pelagic biomass in photic nearshore and/or surface areas (1st row of panels in Fig. 6, Scenario A, C and D). Pelagic biomass therefore increases with a reduction in both horizontal and vertical mixing up to the point where mixing becomes insufficient to supply nutrients and counter sinking losses, producing an L-shaped biomass pattern in mixing space (Fig. 2c).

**Figure 6:**
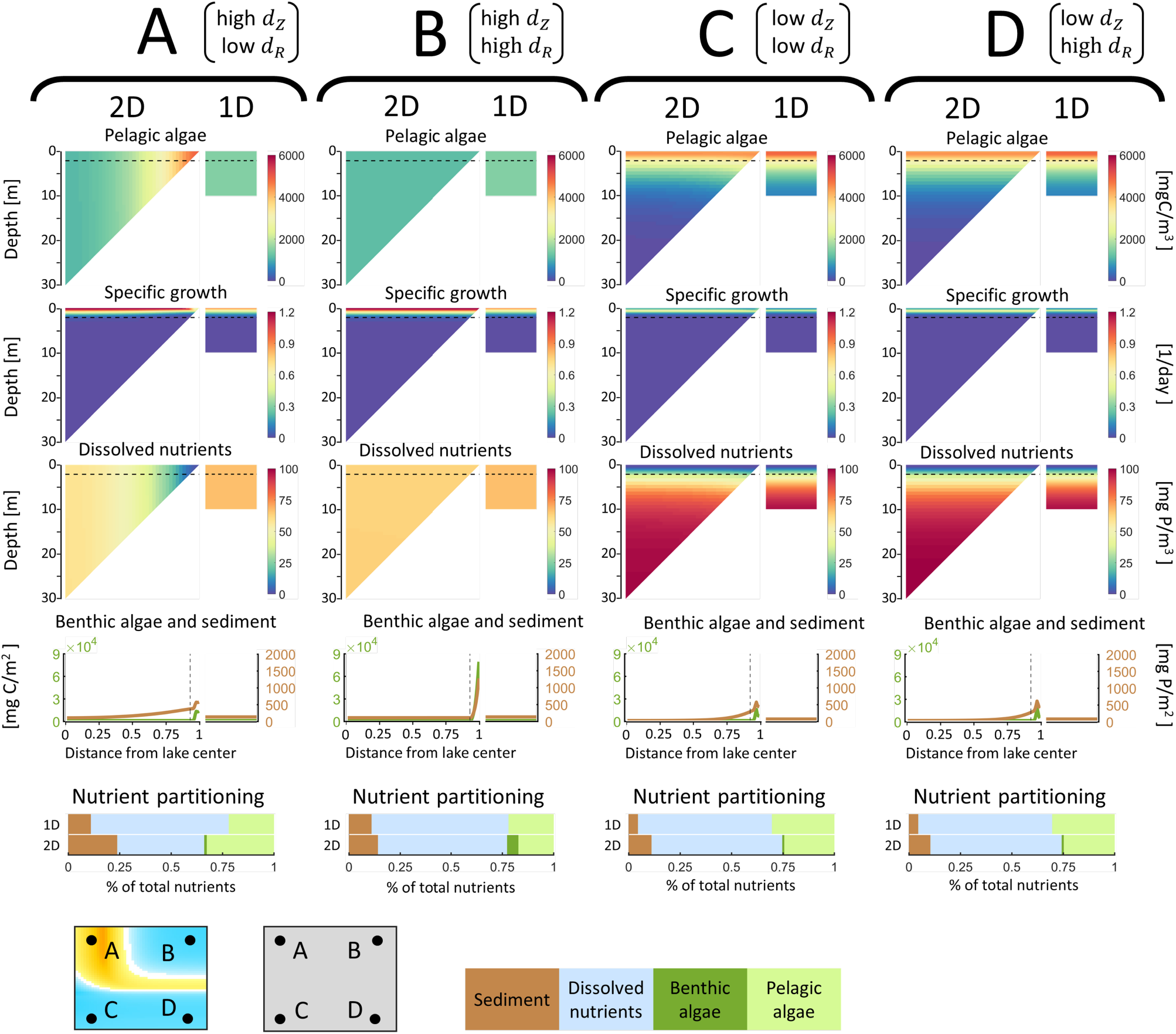
Four example simulations visualizing 2D- and 1D-model output that yields pattern III in Fig. 3. Horizontal broken lines in the first three panel rows mark the boundary of the euphotic depth, below which specific background losses exceed light-dependent production in a hypothetical system lacking pelagic algae. See caption of Fig. 4 for remaining details.

The 1D-model shows the same qualitative pattern along the vertical mixing axis as the 2D-system at high horizontal mixing, but it cannot capture the horizontal pattern since it lacks shallow nearshore areas, yielding an underestimation of pelagic biomass at high vertical and low horizontal mixing (Fig. 2c,f,i, Fig. 6, Scenario A). The 1D-model overestimates pelagic biomass in most of the remaining mixing space, because a smaller fraction of its lake volume is aphotic. The exception is a narrow band of underestimation near *d*_*Z*_ = 10^*−*1^, which arises because the biomass peaks in the vertical mixing direction are slightly offset between the 2D- and 1D-models (Fig. 2c,f,i). Note that in even deeper and darker environments (optical depth = 60), the L-shaped pelagic pattern is amended by yet another type of outcome, i.e. extinction of pelagic algae in both the 2D-and 1D-models (Fig. 3, bottom row of panels, black colored areas). This occurs because 90 % of the lake volume is aphotic. Depth-integrated growth therefore becomes negative at sufficiently high horizontal and vertical mixing.

Finally, the 1D-model again incorrectly predicts the extinction of benthic algae (Fig. 2o, r), because the bottom of the 1D-lake is entirely aphotic, while benthic algae can persist in the shallow margins of the 2D-lake (4^th^ row of panels in Fig. 6). Interestingly, in 2D-mixing space, benthic biomass shows the opposite pattern as pelagic biomass, i.e. an L-shaped pattern with highest biomass at high horizontal and vertical mixing (Fig. 2l). This arises because nutrient supply to nearshore areas is highest in this mixing regime (scenario B in Fig. 6).

In summary, pattern III is similar to pattern II in that the 1D-model incorrectly predicts the extinction of benthic algae, and overestimates pelagic algal biomass at both low vertical mixing and high overall mixing (see ‘Nutrient partitioning’ in Fig. 6). It differs from pattern II primarily in that the 1D-model underestimates pelagic algal biomass in an L-shaped mixing space peaking at high vertical but low horizontal mixing. Note, however, that the transition between these two patterns is not discrete and that weak L-shaped patterns can be sensed in panels in Fig. 3 that we classified as pattern II.

### Robustness of results against inclusion of a mixed surface layer

We evaluated the robustness of the results against the inclusion of a well-mixed surface layer by repeating all simulations across the full 5-dimensional environmental parameter space using two model variants that included a well-mixed surface layer of 1.5 and 4.5 m depth, respectively. Under the vast majority of environmental conditions, results from the three models are in very good qualitative (and often quantitative) agreement with respect to the partitioning of total nutrients among pelagic and benthic algal biomass and dissolved and sedimented nutrients. We therefore describe these results only briefly here, focusing on comparisons of 1D-vs. 2D-model output in both presence and absence of mixed surface layers. To illustrate the results we use the archetypical examples of patterns I-III as shown in Fig. 2 and Fig. D1.

Not surprisingly, when vertical mixing in the standard model is high, results in presence or absence of a mixed surface layer are close to indistinguishable (columns A and B in Fig. D1). In contrast, when vertical mixing in the standard model is low (columns C and D in Fig. D1), including a well-mixed surface layer leads to higher pelagic algal biomass and lower dissolved nutrient concentration in more nutrient-rich and/or optically deeper systems (patterns II and III). Yet, the qualitative differences between 1D- and 2D-models remain the same in these two cases, so the results of the standard model are qualitatively robust. The comparison between 1D- and 2D-models differs qualitatively in presence vs. absence of a mixed surface layer only in the rare (and physically unlikely) case when vertical mixing in the standard model is low, the system is shallow and clear, and the mixed layer depth is equal to the total water column depth of the 1D-model but shallower than the maximum depth of the 2D-model. Under those circumstances, a 1D-model with a mixed surface layer over-(rather than under-)estimates pelagic algal biomass and under- (rather than over-)estimates benthic biomass (pattern I, columns C and D, 4.5 m mixed layer in Fig. D1).

## Discussion

In this study, we used a conceptual modeling approach to address the question: To what extent can pelagic and benthic producer dynamics, integrated over the volume and bottom topography of an idealized, inverted cone-shaped lake basin, be captured by models that use lake mean depth as the only morphometrical input variable? Below, we first discuss qualitative similarities and differences between the predictions from our model Lake2D and its 1D-equivalent, as well as novel phenomena emerging in the 2D-model. We subsequently highlight quantitative differences between the models with a special emphasis on the relative roles of competition between benthic and pelagic algae versus environmental drivers in different regions of the investigated 5-dimensional environmental parameter space. We conclude with a short discussion of implications for whole-lake primary production.

### Qualitative comparison of 1D- and 2D-model predictions – pelagic algae

1D water column models predict that pelagic algal biomass should be maximized at moderate water column depths and turbulence (Diehl, 2002; Huisman & Weissing, 1995; Huisman et al., 2002; Jäger et al., 2010), but that pelagic algae should go extinct under three distinct circumstances: (1) at very low turbulence – when slow upward transport of dissolved nutrients and algae cannot compensate for sinking losses of algae and their sequestered nutrients to aphotic depths or the lake bottom (Huisman et al., 1999; Jäger et al., 2010); (2) in very shallow water columns, where high sinking loss rates exceed growth rates (Diehl, 2002; Huisman & Sommeijer, 2002); and (3) under conditions of high turbulence in combination with high optical depth, when the fractional time spent above compensation depth is insufficient to allow for positive population growth (Huisman et al., 1999; Jäger et al., 2010; Yoshiyama et al., 2009). As discussed below, all of these phenomena can also be observed when the horizontal dimension of lake basins is considered, plus several new phenomena not observable in 1D-models.

Thus, our Lake2D model predicts extinction of pelagic algae below a minimum threshold of the vertical mixing rate, the value of which decreases with both nutrient enrichment and lake depth (since growth is an increasing function of nutrient supply and sinking losses are a decreasing function of lake depth). In our numerical analyses, extinction of pelagic algae at low turbulence is therefore only visible in the shallowest and most nutrient depleted scenarios (black and dark gray areas in upper left panels of Fig. 3), but would be observed in all environmental scenarios at even lower vertical mixing rates than shown. Regardless of the level of turbulence, our model also predicts that a minimum lake depth is required for pelagic algae to persist against sinking losses. Yet, for the relatively low algal sinking velocity that we used (0.1 m day^*−*1^), the theoretical mean lake depth required to drive pelagic algae extinct is so unrealistically shallow that we did not include it in the analyses presented in Fig. 3. Considerably higher sinking velocities would be required to make this mode of extinction ecologically relevant (Huisman & Sommeijer, 2002). Finally, our model also predicts extinction of pelagic algae under conditions of high vertical mixing combined with high optical lake depth (bottom two rows in Fig. 3). Interestingly, extinction occurs at a smaller optical depth (OD = 30) in 2D than in 1D, and extinction depends also on horizontal mixing. Thus, in lakes with a mean depth of 30 m, a background attenuation of 1 m^*−*1^, and high vertical and horizontal mixing, pelagic algae go extinct in the 2D-but not the 1D-model (Fig. 3, gray areas in 2^nd^ row of panels from the bottom), most likely because rapid horizontal mixing away from shallow to deep, vertically well-mixed parts of the lake causes stronger light limitation of pelagic production in 2D-systems (74% of the 2D-lake volume is above bottoms that exceed the depth of the 1D-system). Yet, extinction in the 2D-model does not occur at very low horizontal mixing rates – corresponding to lakes with large surface areas – where pelagic algae can persist in the shallower, relatively well-lit lake margins (cf. Scenario A in Fig. 6).

Finally, similar to 1D-models, we find in optically deeper 2D-lakes that there is an optimal vertical turbulence at which pelagic biomass peaks (Fig. 2c, f). This phenomenon arises, because pelagic algae can sustain a high biomass when vertical mixing is adequate to both transport sufficient nutrients to the photic zone and counteract algal sinking; in contrast, pelagic algae sink out of the photic zone when vertical mixing is too low, and get mixed out of the photic zone when vertical mixing is too high (Huisman et al., 2002; Jäger et al., 2010). Interestingly, pelagic biomass peaks at a higher vertical mixing rate in 2D versus 1D (Fig. 2c, f), likely because, at lower turbulence, a larger fraction of nutrients becomes locked up in nearshore sediments (where pelagic sinking losses are high) and in dissolved form in deeper, aphotic parts of the lake (compare 2D- and 1D-output in scenarios C and D in Fig. 6).

Lake2D reveals also an entirely novel phenomenon – under certain environmental circumstances, there exists an optimal horizontal mixing rate at which pelagic biomass peaks. Specifically, in optically deeper lakes with medium to high vertical mixing, pelagic algae reach a biomass peak at sufficiently low horizontal mixing (Fig. 2c), corresponding to lakes with large surface areas. This biomass maximum is caused by the same mechanism discussed above that prevents pelagic algae from going extinct at even higher optical depths; i.e., horizontal mixing is low enough to prevent pelagic algae in shallow nearshore areas from being mixed to deeper, aphotic parts of the lake but high enough to transport sufficient nutrients from the lake center to pelagic algae in well-lit nearshore areas (Scenario A in Fig. 6).

### Qualitative comparison of 1D- and 2D-model predictions – benthic algae

While 1D- and 2D-models make relatively similar qualitative predictions for pelagic algae, this is not the case for benthic algae. For example, 1D-models predict that benthic biomass decreases monotonically with decreasing light supply (higher optical depth) and goes extinct once a critical minimum threshold of light supply is reached (Jäger & Diehl, 2014; Vasconcelos et al., 2016). None of this is necessarily the case when the horizontal dimension of a lake basin is taken into account. Thus, in the 2D-model benthic algae never go extinct, because they can always persist in sufficiently well lit, shallow nearshore areas. While this is an obvious consequence of including a gradually sloping bottom topography in the model, a less expected result is that benthic biomass can increase with increasing optical depth in the 2D-model. In a comparison of the pattern II and III examples for relatively high vertical and horizontal mixing (Fig. 2k, l and Scenario B in Fig. 5 and 6), benthic biomass approximately doubles as optical depth increases from 3 to 20, in spite of the nutrient loading being the same in both cases. The underlying mechanism is that, at higher optical depths, horizontally and vertically well-mixed pelagic algae become increasingly mixed to aphotic depths and decrease in biomass (Fig. 2b, c), thus liberating nutrients that are transported to well-lit nearshore areas where they support an increasing benthic algal biomass. In contrast, while at high vertical mixing pelagic algae decline with increasing optical depth also in the 1D-model (Fig. 2e, f), benthic algae cannot benefit from the liberated nutrients because the entire 1D-lake bottom is aphotic (Fig. 2n, o).

Lake2D predicts also an entirely novel phenomenon not observable in a 1D context – the potential existence of a benthic algal biomass maximum at some intermediate depth between the shoreline and the lake center (e.g. scenarios C and D in Fig. 5). In analogy to a similar, well-documented phenomenon in the pelagic habitat (Leach et al., 2018), we label this a “benthic deep chlorophyll maximum” (abbreviated to benthic “DCM”). Two conditions are necessary for its occurrence: (i) algal growth must be nutrient limited near the surface and light limited at greater depth; (ii) mineral nutrient concentration must increase with depth. When these conditions are met, nutrient limited specific algal growth will increase with depth until a point of maximum growth is reached where light becomes limiting and growth starts to decrease again towards greater depths with decreasing light. A benthic DCM will then arise at exactly the depth where algal growth switches from nutrient to light limitation.

Since high turbulence prevents the establishment of a vertical gradient in dissolved nutrients, a benthic DCM cannot arise under high vertical mixing (scenarios A and B in Fig. 4-6), but is commonly observed at low vertical mixing rates – as long as a switch from nutrient to light limitation occurs somewhere along the depth gradient (scenarios C and D in Fig. 4-6). Note that the conditions for the establishment of a benthic DCM are identical to the ones for the establishment of a pelagic DCM (Klausmeier & Litchman, 2001; Yoshiyama et al., 2009). In fact, since we assumed identical traits for benthic and pelagic algae, benthic and pelagic DCMs tend to occur in conjunction and at almost the same depth, the pelagic DCM being only slightly offset downwards because of algal sinking (e.g. scenarios C and D in Fig. 4-5).

To our knowledge, the possibility of a benthic DCM, driven by opposing vertical gradients in light vs. nutrient supply, has not yet been proposed. While benthic algal production and biomass are rarely measured along depth transects within lakes, several studies suggest that benthic primary production and chlorophyll a do frequently not peak in the shallowest nearshore areas of lakes but at some distance and depth from the shoreline (Devlin et al., 2016; Gushulak et al., 2021; Klaus et al., 2022; Stevenson & Stoermer, 1981; Vadeboncoeur et al., 2014). A similar pattern has been documented for submerged macrophytes, which typically show a biomass maximum at some intermediate depth, with the depth of this maximum being positively related to water transparency (Duarte & Kalff, 1990). While the latter observation would be in line with the proposed mechanism producing a benthic DCM, the authors of the cited studies unanimously invoked physical disturbance by waves or ice scour as the cause of declining primary production and biomass towards nearshore areas. In contrast, the mechanism promoting a benthic DCM, i.e. increasing nutrient limitation towards shallower depths, has so far not been considered because epipelic algae are believed to have direct access to abundant sediment nutrients (Genkai-Kato et al., 2012; Vadeboncoeur et al., 2008). Yet, both theoretical and empirical studies indicate that benthic primary production can be nutrient limited in oligotrophic lakes (Fork et al., 2020; Hansson, 1992; Vasconcelos et al., 2018). We therefore believe that the possibility of a benthic DCM, driven by opposing vertical gradients in light vs. nutrient supply, deserves further scrutiny.

### Quantitative comparison of 1D- and 2D-model predictions: Roles of competitive interactions vs. environmental drivers

A striking outcome of our analyses is that – across the extensive, 5-dimensional environmental parameter space that we explored – the 1D-model rarely gives similar quantitative results as the 2D-model. Most commonly, the 1D-model underestimates the biomass of benthic algae and overestimates the biomass of pelagic algae. Yet, the reverse can be true in nutrient poor, optically shallow systems, but also in optically deep, vertically well-mixed systems with low horizontal mixing, where pelagic algae can maintain high biomass in shallow nearshore areas that do not exist in a corresponding 1D-model. Note that these results are robust against the relaxation of a potentially problematic feature of our approach, i.e. the assumption that the vertical mixing rate is homogenous over depth. While many of the world’s lakes are seasonally or permanently stratified into a well-mixed surface layer and a much more weakly mixed hypolimnion, our assumption of homogeneous mixing reduces model complexity, which greatly facilitates the interpretation of model output and, moreover, makes results directly comparable to previous work in 1D that also assumed homogeneous vertical mixing (Huisman et al., 2002; Jäger & Diehl, 2014; Jäger et al., 2010). Since simulations with or without mixed layers of two different depths give very similar results (see Fig. D1), we are confident that our conclusions drawn from simulations assuming homogeneous vertical mixing are broadly relevant also for stratified lakes.

Interestingly, our comprehensive exploration reveals the existence of only three archetypical patterns of differences between the 1D- and 2D-models, which are characterized by, respectively, diagonal, vertical or L-shaped gradients of the 1D:2D pelagic biomass quotas in mixing space (Fig. 3). Which of these three patterns arises where in parameter space can be largely explained by the relative importance of competitive interactions between benthic and pelagic algae vs. environmental constraints on the underwater light environment. Specifically, the diagonal pattern, which occurs in nutrient poor, optically shallow systems, is primarily driven by interspecific competition, while the vertical pattern is driven by more pronounced light limitation and the L-shaped pattern by an extremely poor underwater light environment.

To understand this, we note that competition for light between benthic and pelagic algae is inherently asymmetric: while shading from pelagic algae always reduces the light supply to benthic algae, benthic algae cannot directly affect the light supply to pelagic algae. In contrast, competition for nutrients is mutual, its outcome depending on lake depth and turbulent mixing. Thus, in more shallow and/or less turbulent systems, where pelagic sinking losses are relatively high, benthic algae can sequester a large share of the system’s total nutrients and intercept the upwards transport of mineralized nutrients from the sediment to pelagic algae. Conversely, in deeper and more turbulent systems, pelagic algae can sequester and maintain a large share of the system’s total nutrients in more central parts of the lake, thus shading out large parts of the lake bottom and limiting the nutrient supply to benthic algae in the less shaded lake margins.

The consequences of these interaction asymmetries for the impact of competition vs. abiotic drivers on algal performance can be visualized by comparing model runs where each competitor is alone vs. in sympatry with the other. These comparisons provide several insights. First, in line with empirical observations (Hansson, 1988, 1992; Vadeboncoeur et al., 2003), benthic algal biomass is always negatively affected by the presence of pelagic algae (compare Fig. 2j-l with Fig. 7j-l, keeping in mind the different color scales). Second, also in line with empirical observations (Hansson, 1988; Vadeboncoeur et al., 2003), if nutrients are the most limiting resource, i.e. in systems that are nutrient poor and optically shallow (pattern I), pelagic algal biomass is negatively affected by the presence of benthic algae (compare Fig. 2a with Fig. 7a). Third, while the latter applies to both 1D- and 2D-models, the strength of competitive effects differs in horizontal mixing space as described in section. This produces complementary diagonal patterns, where the 1D-model overestimates the competitive effect of benthic algae at low vertical and high horizontal mixing rates, but underestimates their competitive effect under opposite conditions (lower right vs. upper left corners in Fig. 2g, p and in most panels on green background in Fig. 3). Finally, if light is the most limiting resource (patterns II and III), pelagic algal biomass is hardly affected by the presence of benthic algae (compare Fig. 2b, c with Fig. 7b, c).

**Figure 7:**
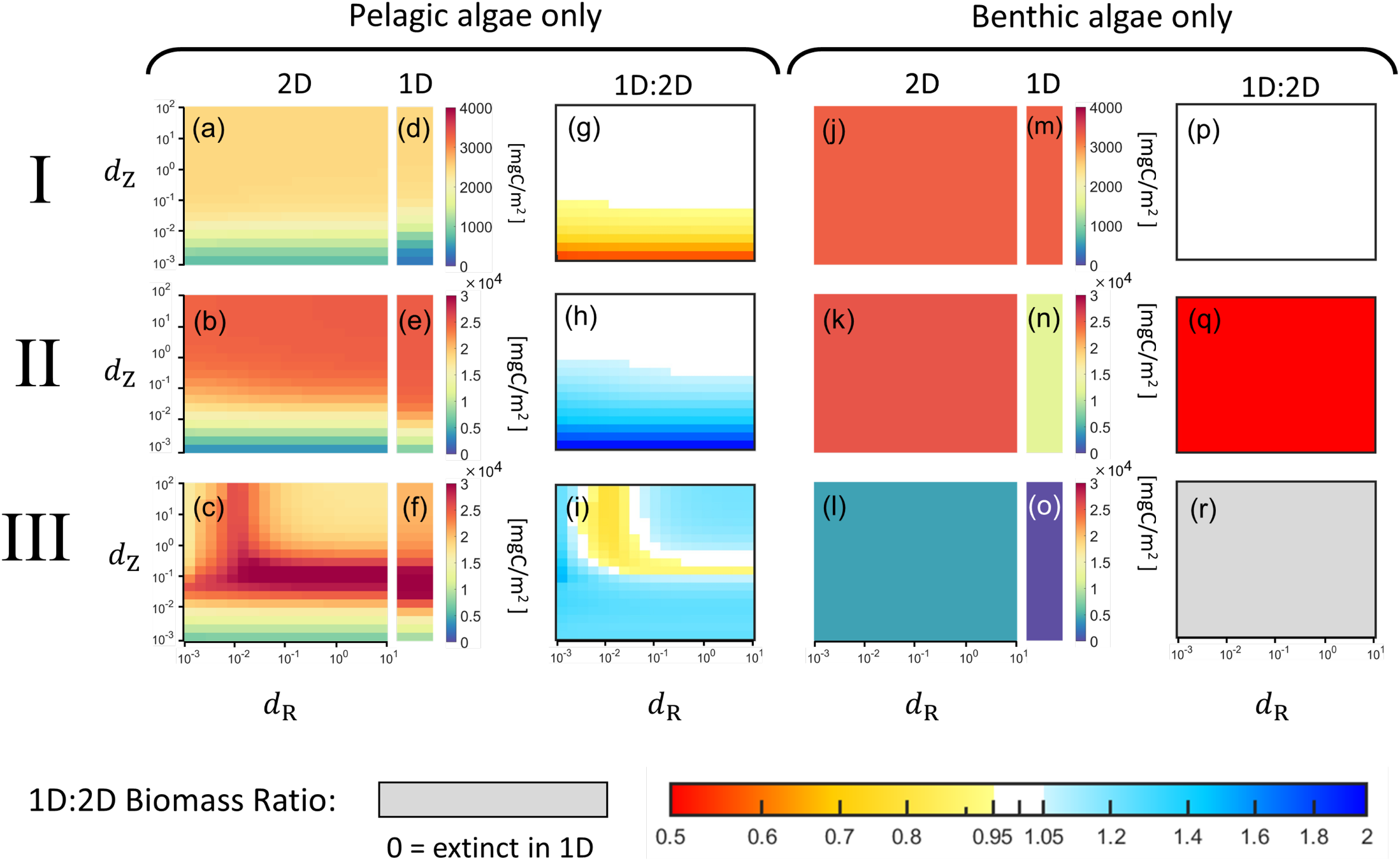
Heat map plots of pelagic and benthic algae in vertical (d_Z_) and horizontal (d_R_) mixing space, where pelagic and benthic algae are simulated in isolation. Panels (a)-(i) show simulations where only pelagic algae are present, and panels (j)-(r) show simulations where only benthic algae are present. Environmental parameters are identical to patterns I-III in Fig. 2. Other parameter values are identical to the original simulations.

The latter observation points to yet another important insight: the observed differences in benthic and pelagic biomass between the 1D- and 2D-models do not represent a zero-sum game between benthic and pelagic algae. This can be illustrated by comparing total algal biomass, i.e. the sum of benthic plus pelagic biomass, between the 1D- and 2D-models. In the larger part of the simulated 5-dimensional parameter space, total biomass differs between the 1D- and 2D-models (Fig. E1), with the 1D:2D ratio of total biomass showing similar patterns as the biomass ratio of the stronger competitor under given environmental conditions. Thus, in clear, shallow systems (largely corresponding to pattern I), total biomass shows similar patterns as benthic biomass, whereas in deeper, more nutrient rich systems, total biomass shows similar patterns as pelagic biomass (compare Fig. 3 with Fig. E1).

In our models, equilibrium algal biomass scales almost linearly with gross primary production (GPP), because we assumed that specific algal losses through respiration and mortality are constant while pelagic algal sinking losses (which vary in space depending on distance to the lake bottom) are small in all but the shallowest lakes. Our results on total algal biomass (Fig. E1) therefore suggest that whole-lake primary production estimates from 1D-models are biased in systematic, predictable ways. Specifically, under the most environmentally relevant mixing scenarios (i.e. high vertical mixing in shallow lakes and lower vertical mixing in deep lakes), the 1D-model tends to underestimate whole-lake GPP by 10-25% (and up to *>* 100%) in both optically shallow lakes corresponding to pattern I and optically deep lakes corresponding to pattern III. Conversely, in lakes of intermediate physical and optical depth, the 1D-model frequently overestimates whole-lake GPP by 10-20%.

## Conclusions

Lakes come in a large variety of different sizes, depths and shapes with shorelines that range from almost circular to highly convoluted (Håkanson, 2004; Scheffers & Kelletat, 2016). In the present analysis, we only considered the two most basic aspects of lake morphometry, i.e. the uneven depth distribution of both the volume and the bottom surface, using an idealized model lake with a radially symmetric geometry. Many ecologically relevant, morphometry-dependent differences between real lakes and our 1D-model approximation, such as fractal shorelines, multi-basin topographies, and the uneven volumetric distribution of temperature across different depth strata (Casas-Ruiz et al., 2021; Puts et al., 2022b; Winslow et al., 2014), were not considered in our study. Despite this simplifying, conceptual approach, we identified several process-based mechanisms that can plausibly explain the empirical observation that lake mean depth is almost always a rather poor predictor of the lake-wide, average production and biomass of benthic primary producers, and frequently also of pelagic and total algal biomass (Carpenter, 1983; Duarte & Kalff, 1990; Puts et al., 2022b).

This being said, our extensive simulations across the 5-dimensional environmental parameter space expose a hierarchy of drivers of lake primary production of which basin morphometry is not the strongest one. As will be described in a separate paper, areal nutrient supply (*N*_area_) has typically the strongest effect by far on whole-lake algal biomass and primary production in the Lake2D model, followed by lake mean depth (through its effects on sinking losses in very shallow lakes and the underwater light climate in deep lakes). Yet, current global estimates of lake GPP neglect both benthic producers and any morphometric information beyond lake area and mean depth (Gao et al., 2021; Jia et al., 2024; Lewis Jr, 2011). Given that our analysis suggests that 1D-approaches underestimate GPP in shallow lakes on the order of 20% and that shallow lakes, because of their shear abundance (Cael et al., 2017) and high productivity (Qin et al., 2020), account for more than half of the estimated global rate of GPP from all freshwater lentic and lotic ecosystems (Rabaey et al., 2024), we believe that global estimates of lake GPP may have to be corrected for the systematic errors inflicted by the prevailing 1D-approaches.

## Supplementary Material

### A Three-dimensional model

Here we describe the full algal model, in three spatial dimensions, from which the 1D and Lake2D model presented in this paper are derived.

Let Ω ⊂ ℝ^3^ be a bounded lake domain with piecewise smooth boundary *∂*Ω equipped with an outward pointing unit normal 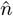. The boundary is partitioned into the lake bottom Γ and the lake surface *∂*Ω \ Γ. We assume the lake surface to be flat and lie in the plane *z* = 0, where *z* is the vertical coordinate with positive direction downwards. The time domain starts at *t* = 0 and ends at a final time *t* = *T*.

The model is comprised of four state variables; pelagic algae *A* and dissolved inorganic nutrient *N*_d_, which are fields within the lake Ω; benthic algae *B* and sedimented particular nutrient *N*_s_, which are fields on the lake bottom Γ. The evolution of these state variables is governed by the system of partial differential equations

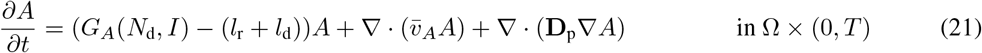

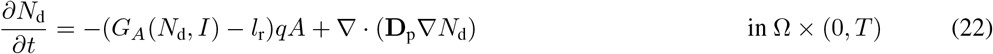

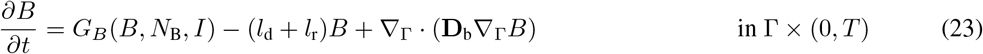

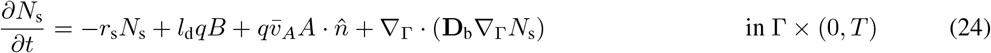

where 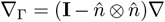 denotes the tangential gradient along the lake bottom 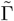, and 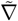 is the usual gradient operator in ℝ^3^. The forms *I, G*_*A*_, and *G*_*B*_ are given by

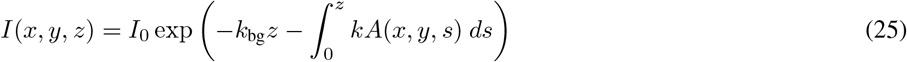

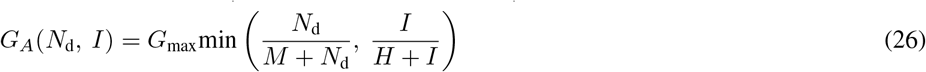

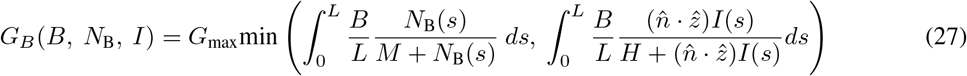

and stems from modeling assumptions described in Section **One-dimensional model**. The parameters in the model are listed in Table 1 in the main text, with the exception that the diffusion coefficients for pelagic and benthic algae are here described as the 3 *×* 3 matrices **D**_p_ and **D**_b_, respectively. The precise structure of these matrices depends on both modeling assumptions and the coordinate system of choice, but in its simplest form with Cartesian coordinates they read **D**_p_ = **D**_b_ = diag([*d*_*x*_, *d*_*y*_, *d*_*z*_]). The unit vector 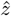 points straight downwards and the unit vector 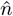 is the normal vector to the lake bottom, pointing outwards from the lake.

#### Benthic dissolved nutrients

The concentration of nutrient concentration in the benthic algal layer is given by the solution to

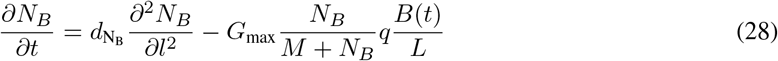

#### Surface boundary conditions

The surface boundary condition for the pelagic algae reads

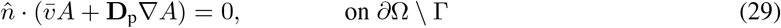

and can be interpreted as zero flux of algae across the lake surface. The boundary condition for the dissolved nutrients reads

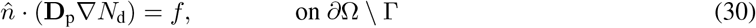

where the function *f* (*x, y, t*) is the outward flux of nutrients across the lake surface. In this article we assume that *f* = 0. A negative value of *f* corresponds to external nutrients flowing into the lake, and a positive value corresponds to dissolved nutrients flowing out of the lake through the surface.

#### Bottom boundary conditions

Pelagic algae sink out at the bottom of the lake, but they do not diffuse out through the lake bottom. This yields the following boundary condition for the pelagic algae:

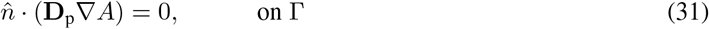

The dissolved nutrients at the bottom are increased by sedimented nutrients remineralizing into the lake and benthic algae dying, releasing their nutrients into the water above. The dissolved nutrients are decreased by consumption from benthic algal growth. This yields the following boundary condition:

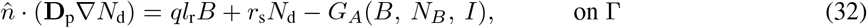

#### Initial conditions

The initial conditions used were as follows: 98% of total nutrients were in dissolved form in the water, 1% of nutrients in pelagic algae, and the remaining 1% in benthic algae. All state variables were well mixed at *t* = 0. The total nutrient content in the system is a fixed quantity with time, due to the choice of boundary conditions and the model formulation (no nutrient sinks or sources inside the lake, or on the lake boundary).

### B Non-dimensionalized system

The non-dimensionalized 3-dimensional system is given by

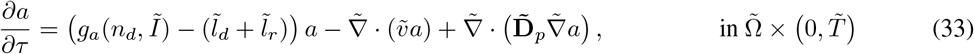

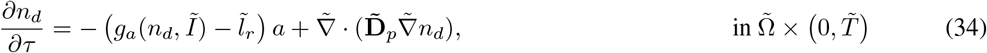

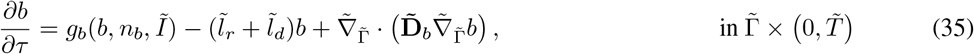

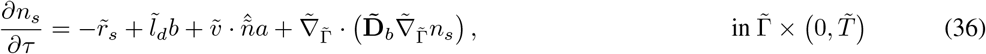

where 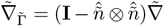 denotes the tangential gradient along the lake bottom 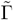, and 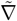 is the usual gradient operator in the non-dimensionalized space. Lowercase letters and tildes are systematically used to indicate the non-dimensionalized forms of the original parameters, state variables, vectors, and operators. The forms *I, g*_*a*_, and *g*_*b*_ are given by

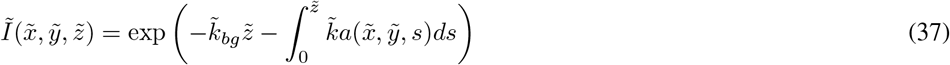

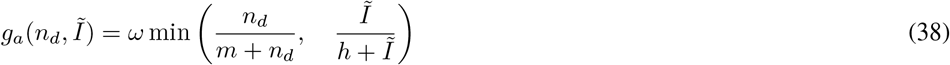

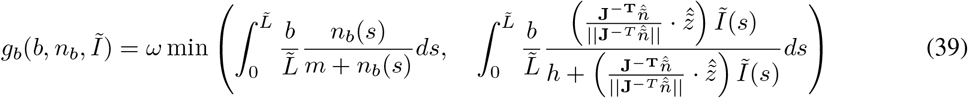

with lake surface boundary conditions

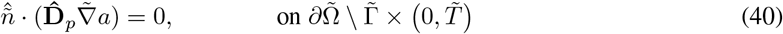

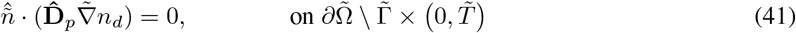

and lake bottom boundary conditions

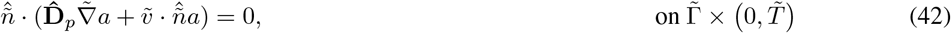

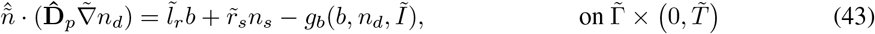

The non-dimensionalized equation for the dissolved nutrients inside the benthic algal layer is given by

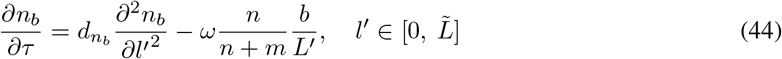

with the boundary condition

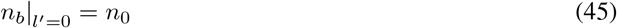

on the surface of the benthic algal layer and

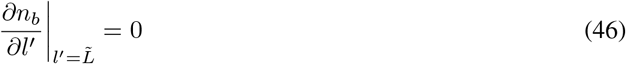

on the bottom of the benthic algal layer. Non-dimensionalized variables and parameters are listed in Table B1 and their definitions are given by

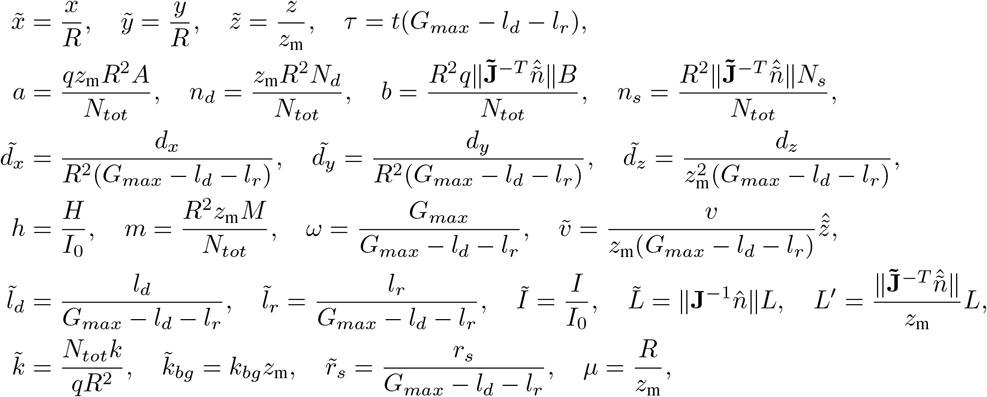

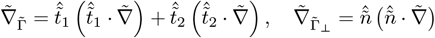, where 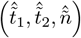 is a basis on the lake bottom, where 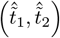 spans the bottom surface and 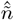 is the lake bottom normal vector.

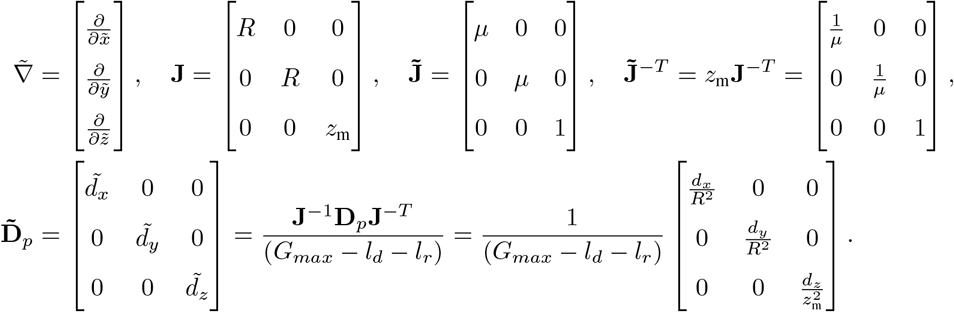

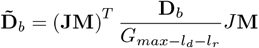, where 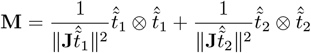, the operator ⊗ is the outer product defined as 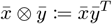 for two arbitrary column vectors 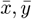.

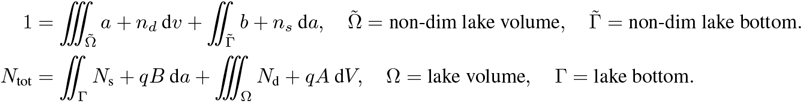

**Table B1:**
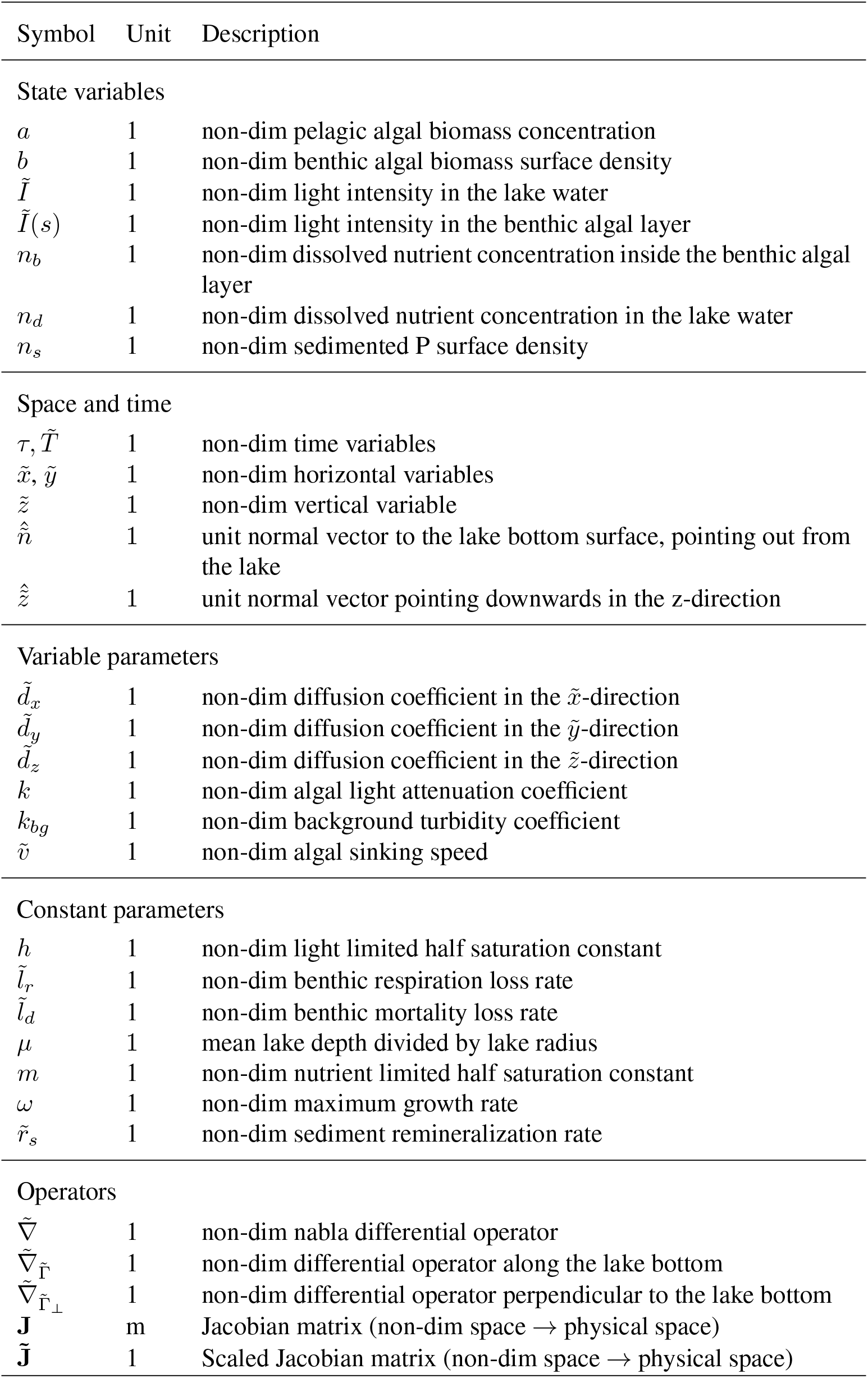
Non-dimensionalized state variables, parameters, operators, and vectors.

### C Numerical method

The finite volume method was used to implement the models presented in this paper. In brief, it involves dividing the lake volume into volume elements (hence the name) and equating the change of biomass or nutrients in each volume element to (1) the fluxes in and out of its edges, and (2) any sources or sinks inside the volume element. This approach has several notable strengths:

- **Nutrient conservation** The finite volume method inherently conserves nutrients. This is because the outward flux from any volume element is always balanced by the inward flux from its neighbor. Consequently, when appropriate boundary conditions are applied to prevent fluxes across the lake’s edges, the total nutrient content within the lake remains constant.
- **Variable diffusion coefficients** A key strength of the finite volume method is its ease of implementation for spatially and temporally variable diffusion coefficients and sinking speeds. The mathematical treatment of these terms remains independent of their variation in space and time, allowing for straightforward modifications. For instance, changing the vertical diffusion coefficient only requires a direct adjustment of the corresponding parameter.
- **Physical interpretation of terms** The finite volume method allows for the physical interpretation of terms in the numerical method. For example, the change of pelagic algal concentration in each volume element is expressed as the sum of algal sinks and sources inside the volume element, i.e. growth and death, and the fluxes in and out across its edges due to sinking and diffusion. This is particularly useful for interpreting the boundary conditions, which define the flux across the lake’s edges.

### D Well-mixed surface layer

In this section, we present simulations analogous to those presented in Fig. 4-6 where we have added a well-mixed surface layer, depicted in Fig. D1. High mixing is achieved by prescribing a large vertical diffusion coefficient from the surface down to a chosen thermocline depth. The horizontal diffusion is identical to the unstratified simulations, as well as the vertical diffusion below the thermocline.

### E Total biomass

In this section we present 1D:2D total biomass quotas of lake-wide total (pelagic + benthic) biomass across 5-parameter space, depicted in Fig. E1. These heatmaps are analogous to the pelagic biomass and benthic biomass 1D:2D quota heatmaps depicted in Fig. 3, using the same simulations and the panels are arranged in the same order.

**Figure D1:**
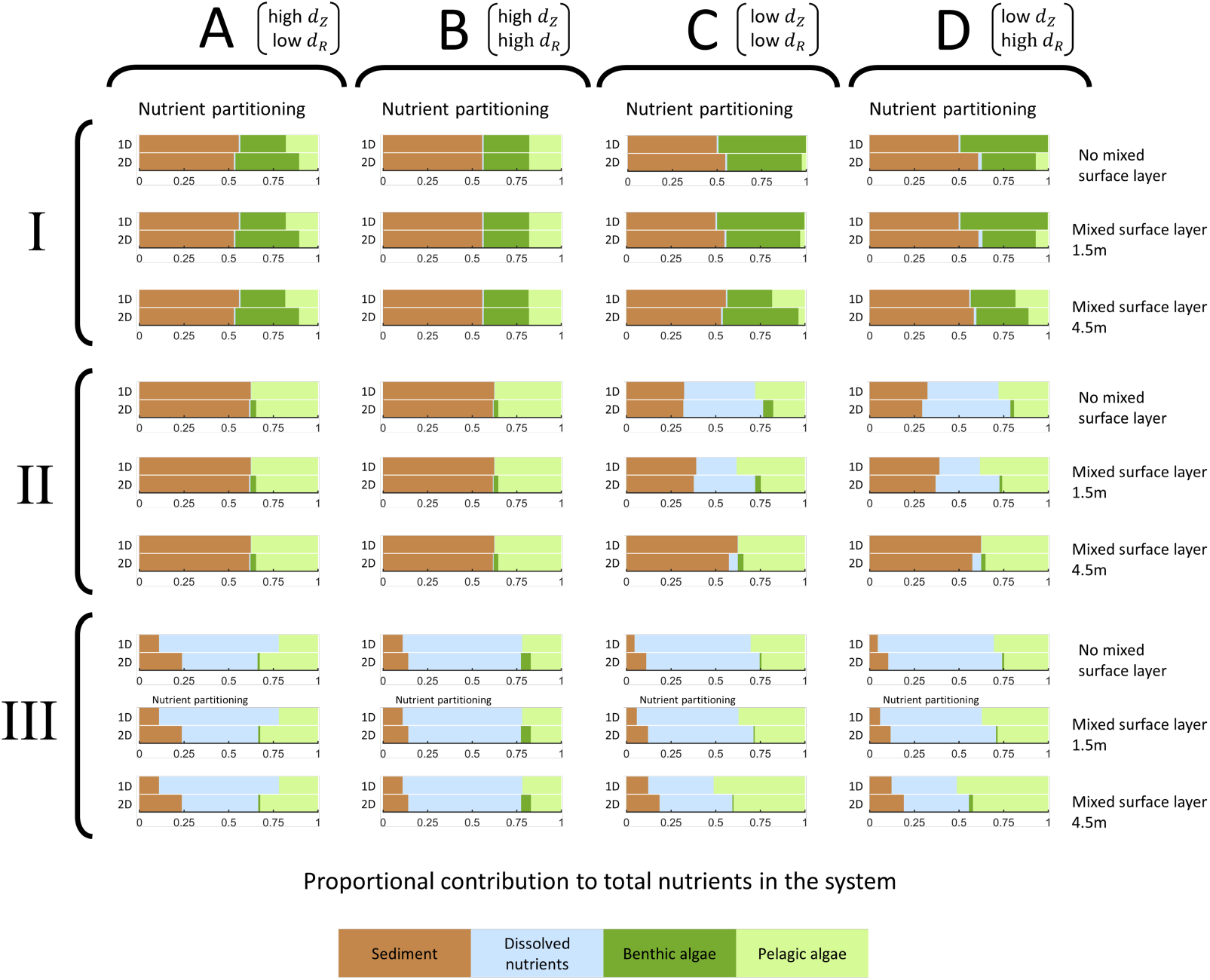
Partitioning of the system’s total nutrients among the four state variables sediment, dissolved nutrients, and benthic and pelagic algae in the standard model with no mixed surface layer and in model runs assuming a well-mixed surface layer of 1.5 m and 4.5 m depth, respectively. Environmental parameter values correspond to patterns I-III (see Fig. 3). Columns indicate the combinations of vertical and horizontal mixing rates in either the entire water body (standard model) or below the mixed surface layer. Column labels A-D correspond to the four corners of mixing space depicted in Fig. 4-6. Each pair of bars shows 1D-output on top and 2D-output below. The portion of total nutrients bound in the sediment is colored brown, dissolved nutrients light blue, benthic algae dark green, and pelagic algae light green.

**Figure E1:**
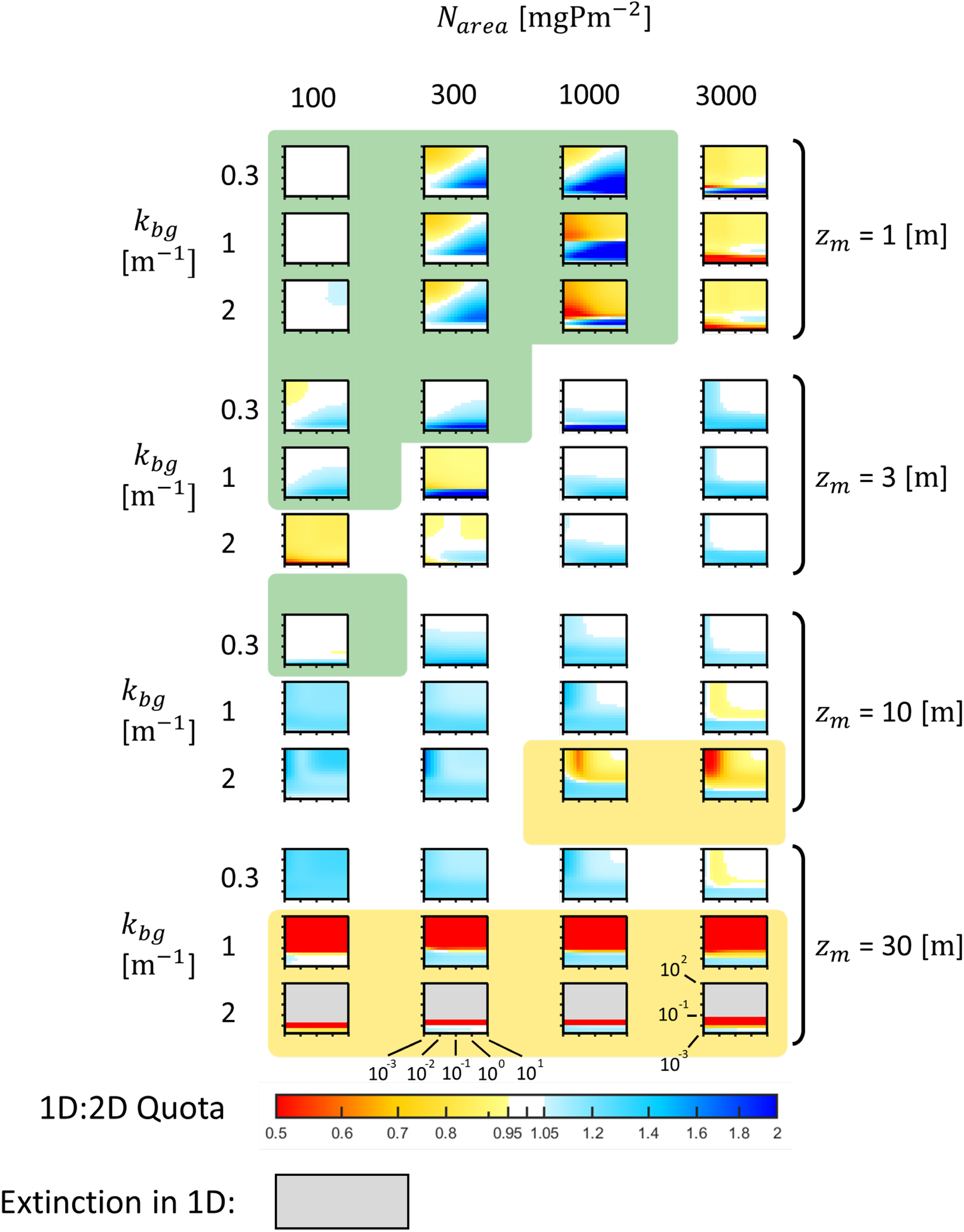
Heat map pairs of 1D:2D quotas of lake-wide total (= pelagic + benthic) algal biomass for varying background turbidity kbg, total nutrient content per surface area N_area_, mean depth z_m_, and diffusive mixing rates d_R_ and d_Z_. Panels are based on the same simulations and are arranged in exactly the same order as the panels in Fig. 3, which show the corresponding, separate heat maps for the 1D:2D biomass quotas of pelagic and benthic algae.

## Notes

### Competing Interest Statement

The authors have declared no competing interest.

